# USP11 deubiquitinates monoubiquitinated SPRTN to repair DNA-protein crosslinks

**DOI:** 10.1101/2020.06.30.180471

**Authors:** Megan Perry, Sai Sundeep Kollala, Meghan Biegert, Grace Su, Manohar Kodavati, Halle Mallard, Natasha Kreiling, Alexander Holbrook, Gargi Ghosal

## Abstract

DNA-protein crosslinks (DPCs) are toxic DNA lesions that interfere with DNA metabolic processes such as replication, transcription and recombination. SPRTN is a replication-coupled DNA-dependent metalloprotease that cleaves proteins crosslinked to DNA to promote DPC repair. SPRTN function is tightly regulated by a monoubiquitin switch that controls SPRTN chromatin accessibility during DPC repair. The deubiquitinase regulating SPRTN function in DPC repair is unknown. Here, we identify USP11 as a SPRTN deubiquitinase. USP11 interacts with SPRTN and cleaves monoubiquitinated SPRTN in cells and *in vitro.* USP11 depletion impairs SPRTN deubiquitination in response to formaldehyde-induced DPCs. Loss of USP11 causes an accumulation of unrepaired DPCs and cellular hypersensitivity to treatment with DPC-inducing agents. Our findings elucidate the function of USP11 in the regulation of SPRTN monoubiquitination and SPRTN-mediated DPC repair.

## INTRODUCTION

DNA-protein crosslinks (DPCs) are irreversible covalent crosslinking of proteins to the DNA. DPCs can be generated by the action of oxygen free radicals, reactive nitrogen species and reactive aldehydes generated as by-products of cellular respiration and metabolism or by exposure to exogenous DNA damaging agents like UV radiation, ionizing radiation (IR) and chemotherapeutic drugs (Barker et al., 2005, Tretyakova et al., 2015). Covalent crosslinking of proteins to the unperturbed duplex DNA, generated by formaldehyde, IR, UV rays and platinum-based chemotherapeutic drugs are classified as type 1 DPCs. Trapped DNA Polβ and PARP1 at the 5′ and 3′ ends of single-stranded DNA breaks (SSBs), respectively, represent type 2 DPCs. Type 3 and type 4 DPCs arise from abortive topoisomerase-DNA enzymatic reactions, resulting in the crosslinking of topoisomerase I (TOP1) to the 3′ end of a SSB or topoisomerase II (TOP2) to the two 5′ ends of a double-stranded DNA break (DSB) (Ide et al., 2018). Crosslinking of specific DNA metabolizing enzymes, such as TOP1 and TOP2, DNA polymerase, and DNA methyl transferase (DNMT1), are also known as enzymatic DPCs (Stingele et al., 2017). Irrespective of the type or source of the lesion, all DPCs are steric blockades that disrupt DNA replication, transcription, recombination and repair processes. Unrepaired or mis-repaired DPCs lead to genome instability, resulting in tumorigenesis and genetic diseases (Stingele et al., 2017, Fielden et al., 2018).

Given the different types of DPCs and myriad of agents that generate them, the precise molecular mechanism underlying the DPC repair pathway has remained elusive. Genetic and biochemical experiments in different model organisms have suggested the nucleotide excision repair (NER) and homologous recombination (HR) pathways mitigate the genotoxic effects of DPCs (Nakano et al., 2007, Ide et al., 2011, de Graaf et al., 2009, Baker et al., 2007). The repair of TOP1 cleavage complexes (TOP1-ccs) and TOP2 cleavage complexes (TOP2-ccs) has been widely investigated. TOP1-ccs and TOP2-ccs are enzymatic DPCs repaired by tyrosyl-DNA phosphodiesterases TDP1 and TDP2, respectively. TDP1 and TDP2 directly hydrolyze the phosphotyrosyl bond between the protein and DNA in the DPC. Following DPC removal, the DNA breaks are subsequently repaired by SSB, HR or non-homologous end joining repair pathways (Pommier et al., 2014, Ide et al., 2018). The proteasome also aids the removal of TOP1-ccs, TOP2-ccs, Polβ crosslinked to DNA and DPCs generated by formaldehyde. However, polyubiquitination of DPCs was observed only in TOP1-ccs and TOP2-ccs, not in formaldehyde-induced DPCs (Nakano et al., 2009, Quievryn and Zhitkovich, 2000). Studies in *Xenopus* egg extracts showed that when the replisome collides with DPCs, the CMG helicase stalls and the DPC is proteolyzed into a peptide-DNA adduct that is bypassed by translesion synthesis (TLS) polymerases, but proteasome inhibition had no significant effect on DPC repair (Duxin et al., 2014). Concurrently, yeast Wss1 was identified as a DNA-dependent metalloprotease that cleaves both enzymatic TOP1-ccs and non-enzymatic formaldehyde-induced DPCs during S-phase (Stingele et al., 2014). Subsequently, SPRTN protease was shown to repair DPCs in mammalian cells (Stingele et al., 2016, Vaz et al., 2016, Lopez-Mosqueda et al., 2016). Recently, Ddi1 aspartic protease was identified in yeast and was shown to repair DPCs independent of the 20S proteasome (Serbyn et al., 2019). Collectively, these studies suggest that DPCs are degraded and removed by a repair pathway that is dependent on either the proteasome or a specific protease.

SPRTN (also known as DVC1/ C1orf124), the mammalian functional homolog of yeast Wss1, is a replication-coupled DNA-dependent metalloprotease (Stingele et al., 2016, Vaz et al., 2016). SPRTN was initially identified as a protein required for repair of UV-induced DNA lesions, restart of stalled DNA replication forks, and as a regulator of TLS (Ghosal et al., 2012, Centore et al., 2012, Mosbech et al., 2012, Juhasz et al., 2012, Davis et al., 2012, Machida et al., 2012, Kim et al., 2013). SPRTN associates with the DNA replication machinery, and loss of SPRTN impaired replication fork progression (Vaz et al., 2016). SPRTN protease activity is mediated by the SprT domain of SPRTN, which contains the HEXXH catalytic motif. Like Wss1, SPRTN protease cleaves TOP1, TOP2, Histone H1, H2A, H2B, H3 and HMG1 in the presence of single-stranded DNA (ssDNA) (Vaz et al., 2016, Stingele et al., 2016). SPRTN also drives auto-proteolysis *in trans* in the presence of ssDNA and double-stranded DNA (dsDNA) (Li et al., 2019). Crystal structure of the SprT domain revealed a metalloprotease sub-domain and Zn^2+^-binding sub-domain, which regulate ssDNA binding and protease activity of SPRTN (Li et al., 2019). SPRTN depletion sensitized cells to treatment with formaldehyde and etoposide, suggesting a role of SPRTN in the repair of formaldehyde-induced DPCs and TOP2-cc, respectively (Stingele et al., 2016, Vaz et al., 2016, Lopez-Mosqueda et al., 2016). In humans, biallelic mutations in *SPRTN* leads to Ruijs-Aalfs syndrome (RJALS) characterized by genome instability, segmental progeria and early-onset hepatocellular carcinoma. RJALS patient cells were defective in SPRTN protease activity, displayed defects in replication fork progression and hypersensitivity to DPC-inducing agents (Lessel et al., 2014). Loss of *Sprtn* in mice resulted in embryonic lethality, while *Sprtn* hypomorphic mice recapitulated some of the progeroid phenotypes and developed spontaneous tumorigenesis in the liver with increased accumulation of DPCs in the liver tissue. Mouse embryonic fibroblasts from *Sprtn* hypomorphic mice displayed accumulation of unrepaired TOP1-ccs and were hypersensitive to treatment with DPC-inducing agents (Maskey et al., 2014, Maskey et al., 2017). These studies showed that SPRTN metalloprotease repairs replication-coupled DPCs in the genome, thereby protecting cells from DPC-induced genome instability, cancer and aging.

A recent study performed in *Xenopus* egg extracts showed that both SPRTN and the proteasome can repair replication-coupled DPCs but are activated by distinct mechanisms. The recruitment of the proteasome to DPCs required DPC polyubiquitination, while SPRTN was able to degrade non-ubiquitinated DPCs. SPRTN-mediated DPC proteolysis depended on the extension of the nascent DNA strand to within a few nucleotides of the DPC lesion, indicating that polymerase stalling at a DPC on either the leading or lagging strand activates SPRTN. SPRTN depletion impaired TLS following DPC proteolysis in both proteasome-mediated and SPRTN-mediated replication-coupled DPC repair, suggesting that in addition to DPC proteolysis, SPRTN regulates bypass of peptide-DNA adducts by TLS during DNA replication (Larsen et al., 2019).

SPRTN is a sequence-nonspecific protease that predominantly cleaves substrates in unstructured regions in the vicinity of lysine, arginine and serine residues (Vaz et al., 2016, Stingele et al., 2016). Several mechanisms regulate SPRTN function in DPC repair. SPRTN protease activity is stimulated by DNA binding, while post-translational modification of SPRTN governs both its protease activity and recruitment to the DPC on chromatin. CHK1 kinase phosphorylates SPRTN at the C-terminal (S373, S374 and S383) and enhances SPRTN protease activity and recruitment to chromatin (Halder et al., 2019). SPRTN is also monoubiquitinated which prevents SPRTN access to chromatin (Mosbech et al., 2012, Stingele et al., 2016). Upon DPC induction, monoubiquitin on SPRTN is cleaved by a ubiquitin protease, allowing SPRTN to localize to chromatin and cleave DPCs in the presence of ssDNA (Stingele et al., 2016). However, the E3 ubiquitin ligase and deubiquitinase (DUB) that regulate SPRTN monoubiquitination are unknown. Here, we have identified USP11 as a ubiquitin protease that regulates SPRTN monoubiquitination upon DPC induction.

USP11 (Ubiquitin carboxyl-terminal hydrolase or ubiquitin specific protease 11) belongs to the ubiquitin specific protease (USP or UBP) family of DUBs. USP11 participates in processes such as TGFβ signaling, pro-inflammatory signaling, viral replication and NF-κB signaling by regulating the protein stability of various targets such as ARID1A, TβRII, CDKN2A, RAE1, XIAP, HPV16-E7 and IκBα (Luo et al., 2020, Deng et al., 2018, Jacko et al., 2016, Lin et al., 2008, Sun et al., 2010, Stockum et al., 2018). USP11 also functions in DSB repair, wherein USP11 deubiquitinates H2AX to regulate the recruitment of RAD51 and 53BP1 to damage foci (Ting et al., 2019, Yu et al., 2016). USP11 deubiquitinates PALB2 and promotes BRCA1-PALB2-BRCA2 complex formation (Orthwein et al., 2015).

USP11 confers cellular resistance to PARP1 inhibitors that trap PARP1 to DNA (Wiltshire et al., 2010). However, the function of USP11 in DPC repair is not reported. In this study, we show that USP11 is a novel interactor of SPRTN and deubiquitinates SPRTN in cells and *in vitro*. Depletion of USP11 leads to an accumulation of unrepaired DPCs and sensitizes cells to DPC-inducing agents. USP11 cleaves the monoubiquitin on SPRTN upon DPC induction and regulates SPRTN-mediated DPC repair.

## RESULTS

### USP11 is a SPRTN interacting protein

Monoubiquitinated SPRTN is deubiquitinated and the unmodified SPRTN is localized to chromatin upon DNA damage by DPC-inducing agents (Stingele et al., 2016). Therefore, ubiquitination and deubiquitination of SPRTN is critical for the recruitment of SPRTN to DPC lesions. To identify SPRTN modifiers, we performed tandem-affinity purification and mass spectrometry (TAP-MS) analysis of SFB (S-FLAG-streptavidin binding peptide)-tagged SPRTN expressed in HEK 293T cells. Our MS analysis revealed several known SPRTN interactors, namely, PCNA, POLD3 and VCP (**Figure 1A and Supplemental File 1**). We screened the MS list for proteins that are known to function as a deubiquitinase and identified only one potential SPRTN deubiquitinase, USP11. Similar to POLD3, only one peptide of USP11 was immunoprecipitated in SPRTN MS analysis. A reciprocal TAP-MS analysis of SFB-USP11 expressed in HEK 293T cells immunoprecipitated SPRTN in addition to USP7, which is a known USP11 interactor **(Figure 1B and Supplemental File 2)**. We next confirmed SPRTN-USP11 interaction by immunoprecipitation in HEK 293T cells expressing either SFB-USP7, SAMHD1, or SPRTN. USP7 is a known interactor of USP11, while SAMHD1 and SPRTN were identified in SFB-USP11 TAP-MS analysis **(Figure 1B)**. Endogenous USP11 was immunoprecipitated with SFB-USP7 and SPRTN, but not with SAMHD1 **(Figure 1C)** and the interaction between SPRTN and USP11 increased following treatment with etoposide (VP16), a TOP2 crosslinking agent **(Figure 1D)**.

**Figure 1.**
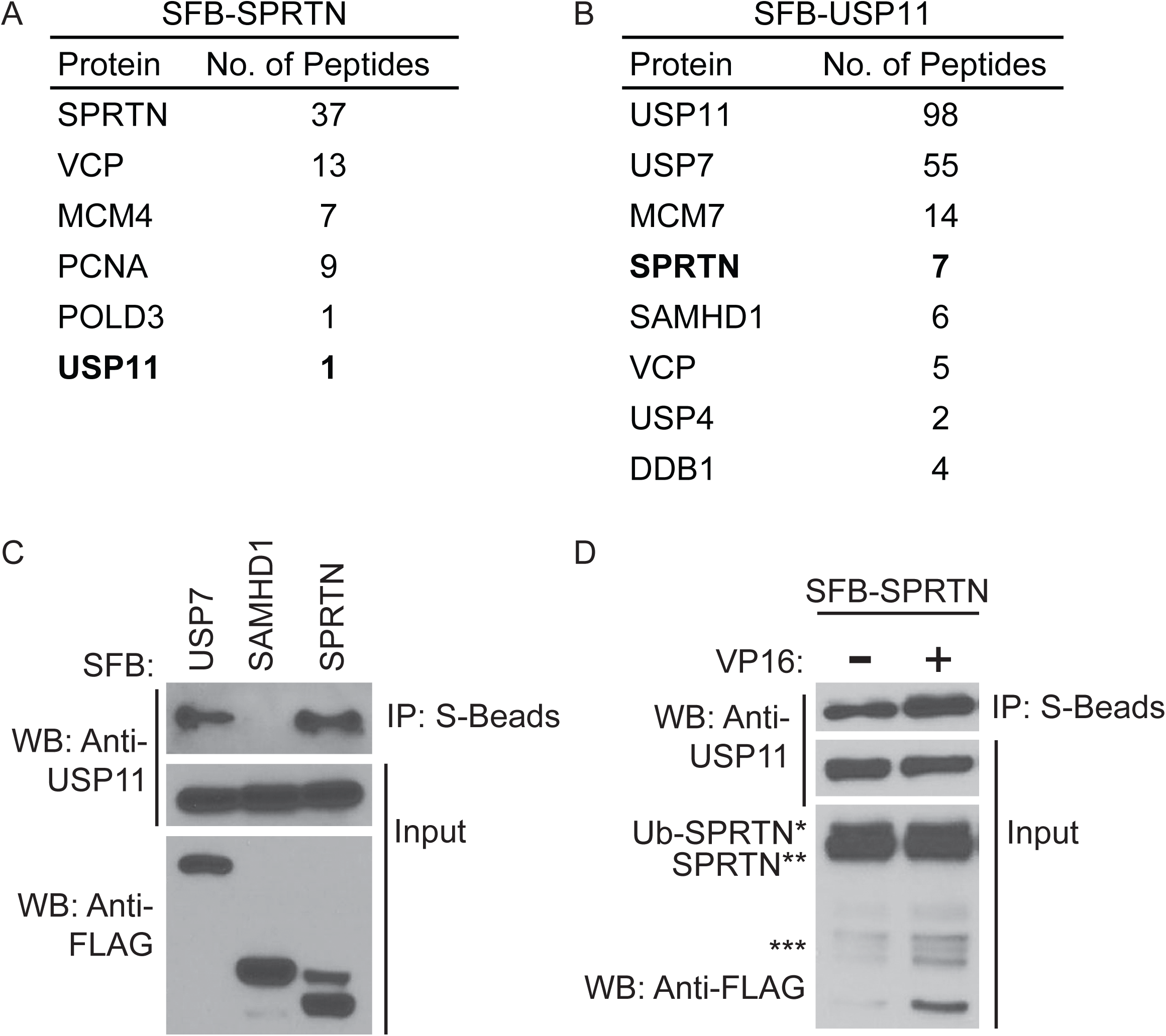
USP11 is a SPRTN interacting protein. (A-B) TAP-MS was performed using HEK 293T cells stably expressing SFB-(A) SPRTN or (B) USP11. SFB-USP11 cells were treated with 10 μM CPT for 2 hr. Selected results from MS analysis (Supplemental File 1 and File 2) are shown. MS data have been deposited to the ProteomeXchange Consortium with the dataset identifier PXD019923. (C) USP11 interacts with SPRTN. HEK 293T cells were transfected with SFB-USP7, SAMHD1, or SPRTN. (D) USP11 interacts with SPRTN in the absence and presence of DPCs. HEK 293T cells transfected with SFB-SPRTN were left untreated or treated with 20 μM VP16 for 4 hr. (C-D) Cell lysates were immunoprecipitated with S-protein agarose beads, and immunoblotting was performed using the indicated antibodies. WB, Western blot. ***indicates SPRTN auto-cleavage products.

Both SPRTN and USP11 are multi-domain proteins. To map interaction sites on SPRTN and USP11, we generated domain deletion mutant constructs of SPRTN (ΔSprT, ΔSH, ΔPIP and ΔUBZ) and USP11 (ΔDUSP and ΔUSP). Co-immunoprecipitation (Co-IP) of Myc-USP11 with SFB-SPRTN full length (FL) or ΔSprT, ΔSH, ΔPIP, ΔUBZ, E112A catalytic inactive and Y117C (SPRTN mutation identified in RJALS patients) mutant constructs showed that SPRTN-USP11 interaction was lost with deletion of the N-terminal SprT domain of SPRTN **(Figure 2A)**. Co-IP experiments of SFB-SPRTN with Myc-USP11 FL, C318S catalytic inactive, ΔDUSP and ΔUSP mutant constructs showed that SPRTN interacts with the C-terminal USP domain of USP11 **(Figure 2B)**. Because the USP domain of USP11 is 654 aa long, we generated several internal deletion constructs within the USP domain. Co-IP experiments of SFB-SPRTN with Myc-USP11 FL and internal deletion constructs of the USP domain showed that the SPRTN interaction site resides within the 480-505 aa region of USP11 **(Figure 2C)**. Similarly, we generated internal deletion mutants within the SprT domain of SPRTN to map the site of USP11 interaction in SPRTN. We found that deletion of the SprT domain and 129-212 aa region in SPRTN abrogated USP11 binding. However, deletion of 186-212 aa fragment in SPRTN retained USP11 interaction, indicating that the USP11 binding site lies within 129-186 aa region of SPRTN (**Figure 2D**).

**Figure 2.**
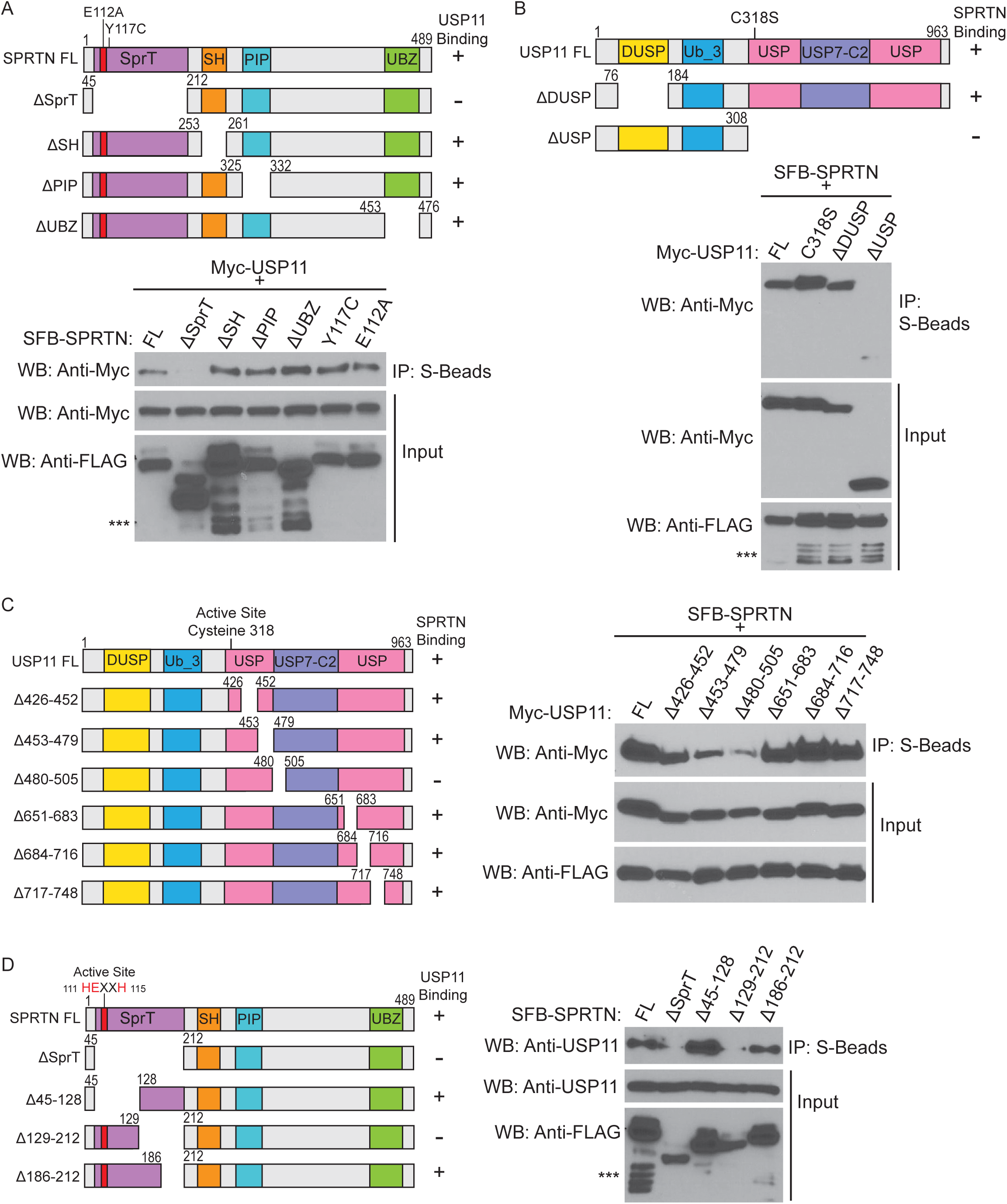
SprT domain of SPRTN and USP domain of USP11 mediate SPRTN-USP11 interaction. (A) Top panel, protein schematic of SPRTN FL, domain deletion and point mutants. Bottom panel, HEK 293T cells were transfected with Myc-USP11 and SFB-SPRTN FL, deletion or point mutants. (B) Top panel, schematic of USP11 FL and deletion mutants. USP11 domains derived from Pfam predicted motifs. Bottom panel, HEK 293T cells were transfected with SFB-SPRTN and Myc-USP11 FL, deletion or point mutants. (C) Left panel, protein schematic of USP11 FL and USP domain deletion mutants. Right panel, HEK 293T cells were transfected with SFB-SPRTN and Myc-USP11 FL or USP domain deletion mutants. (D) Left panel, protein schematic of SPRTN FL and SprT domain deletion mutants. Right panel, HEK 293T cells were transfected with SFB-SPRTN FL and SprT domain deletion mutants. (A-D) Cell lysates were immunoprecipitated with S-protein agarose beads, and immunoblotting was performed using the indicated antibodies. WB, Western blot. ***indicates SPRTN auto-cleavage products.

### USP11 deubiquitinates SPRTN in cells and *in vitro*

Based on our observation that SPRTN and USP11 interact **(Figure 1 and 2)** and because USP11 is a deubiquitinase (Luo et al., 2020, Deng et al., 2018, Orthwein et al., 2015), we investigated whether SPRTN is a substrate for USP11 deubiquitinase. Indeed, we found that overexpression of Myc-USP11 FL, but not C318S catalytic inactive mutant was able to deubiquitinate SFB-SPRTN in HEK 293T cells **(Figure 3A)**. Previous studies have shown that SPRTN is monoubiquitinated (Mosbech et al., 2012, Stingele et al., 2016). To confirm monoubiquitination of SFB-SPRTN, we transfected SFB-tagged vector, SPRTN or SPRTN and HA-ubiquitin (HA-Ub) in HEK 293T cells, immunoprecipitated and immunoblotted with FLAG, HA, and ubiquitin (FK2 and K63-Ub) antibodies. We observed that the FK2 antibody, which recognizes both monoubiquitin and polyubiquitin chains, probed the modified band of SPRTN (Ub-SPRTN). Notably, FK2 antibody recognized only a single modified band indicating that the modification on SPRTN corresponds to monoubiquitination (**Figure 3B, anti-Ub FK2 IP blot**). We also found that K63-Ub antibody recognized only the single monoubiquitin band and no K63-linked Ub-chains were observed (**Figure 3B, anti-Ub K63 IP blot**). FK2 and K63-Ub antibodies probed the modification on SPRTN in SFB-SPRTN and SFB-SPRTN-HA-Ub and not in SFB-vector immunoprecipitated samples. Notably, SPRTN modified by HA-Ub was recognized by HA antibody in SFB-SPRTN-HA-Ub, but not in SFB-vector or SFB-SPRTN immunoprecipitations confirming that the modification on SPRTN corresponds to ubiquitination. Similar to FK2 and K63-Ub antibodies, HA-antibody probed a single HA-Ub band and no polyubiquitination was observed (**Figure 3B, anti-HA IP blot**). These observations indicate that the modification on SPRTN corresponds to monoubiquitination.

**Figure 3.**
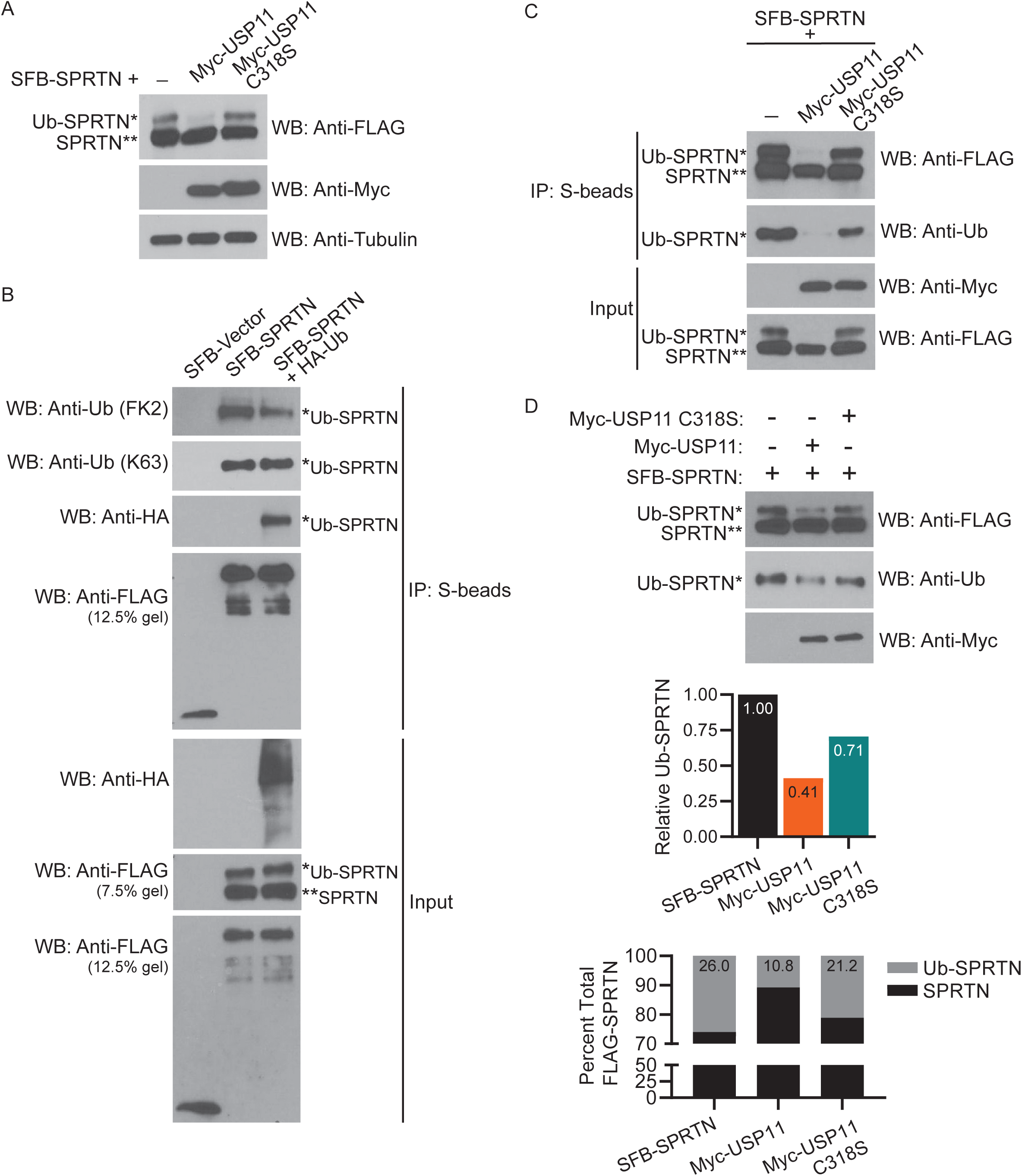
USP11 deubiquitinates SPRTN in cells and *in vitro*. (A) HEK 293T cells were transfected with SFB-SPRTN and Myc-USP11 FL or C318S. Cell lysates were immunoblotted with the indicated antibodies. (B) HEK 293T cells were transfected with SFB-empty vector, SFB-SPRTN, or SFB-SPRTN and HA-Ubiquitin. (C) HEK 293T cells were transfected with SFB-SPRTN and Myc USP11 FL or C318S mutant. (B-C) Cell lysates were immunoprecipitated with S-protein agarose beads and immunoblotted with indicated antibodies. (D) USP11 deubiquitinates SPRTN *in vitro.* Top panel, SFB-SPRTN, Myc-USP11, and Myc-USP11 C318S were purified as described in methods. SFB-SPRTN was incubated with Myc-USP11 or Myc-USP11 C318S mutant proteins overnight at 30 °C. The reaction mixture was immunoblotted with indicated antibodies (C and D, anti-Ub blots were probed with K63 ubiquitin antibody). Middle panel, graph shows relative UbSPRTN values quantified from anti-Ubiquitin WB. Bottom panel, graph shows percent total-FLAG SPRTN. Percent of unmodified and monoubiquitinated SPRTN was calculated from anti-FLAG WB. WB, Western Blot. See also Supplemental Figure 1.

We next confirmed deubiquitination of monoubiquitinated SPRTN by performing deubiquitination assays in HEK 293T cells expressing SFB-SPRTN either alone or in combination with Myc-USP11 FL or C318S mutant (**Figure 3C**) and SFB-SPRTN and HA-Ub constructs expressed either alone or in combination with Myc-USP11 FL and C318S mutant (**Supplemental Figure 1A**). SFB-SPRTN was immunoprecipitated and monoubiquitination of SPRTN was examined by immunoblotting with Ub (**Figure 3C**) or HA antibodies (**Supplemental Figure 1A**). We observed that monoubiquitinated SPRTN is deubiquitinated by USP11 FL, but not C318S deubiquitinase inactive mutant (**Figure 3C and Supplemental Figure 1A**). To show that deubiquitination of SPRTN by USP11 is direct, we purified SFB-SPRTN, Myc-USP11 and Myc-USP11 C318S proteins from HEK 293T cells and performed an *in vitro* DUB reaction by incubating SFB-SPRTN alone or with Myc-USP11 or Myc-USP11 C318S purified proteins. Deubiquitination of SPRTN was examined by immunoblotting with ubiquitin antibody (**Figure 3D**). We observed that monoubiquitination of SPRTN was dramatically reduced in USP11 FL compared to USP11 C318S mutant (**Figure 3D**). We further corroborated this observation in a similar *in vitro* DUB reaction by incubating SFB-HA-UbSPRTN alone or with Myc-USP11 or Myc-USP11 C318S purified proteins and immunoblotted for HA-Ub (**Supplemental Figure 1B**). We observed a marginal reduction in monoubiquitinated SPRTN with USP11 C318S mutant **(Figure 3D and Supplemental Figure 1B)** which could be due to only partial loss of USP11 deubiquitinase activity, as the catalytic core of all UBP family DUBs are comprised of both a Cys (nucleophile) and His (proton acceptor) residue (Hu et al., 2002). Together these findings demonstrate that USP11 deubiquitinates monoubiquitinated SRTN in cells and *in vitro*.

### USP11 is required for cell survival upon DNA damage by DPC-inducing agents

SPRTN is required for repair of DPCs (Vaz et al., 2016, Stingele et al., 2014). However, the function of USP11 in response to DPC lesions and DPC repair is not known. To gain insight into the functional relevance of SPRTN-USP11 interaction, we first examined whether USP11 functions in response to DNA damage caused by DPC-inducing agents. We generated USP11 knockdown in U2OS (**Figure 4A**) and A549 (**Supplemental Figure 2A**) cells to rule out cell type specificity effects in the experiments. Negative control and USP11 knockdown U2OS and A549 cells were untreated or treated with increasing concentrations of DPC-inducing agents, and clonogenic cell survival was examined 10 days post-treatment (**Figure 4 and Supplemental Figure 2**). We found that depletion of USP11 sensitized U2OS cells to treatment with TOP1 crosslinking agent, camptothecin (CPT) **(Figure 4B)**, VP16 **(Figure 4C)**, and non-specific DNA-protein crosslinker, formaldehyde **(Figure 4D)**. We observed mild sensitivity of USP11 knockdown cells to hydroxyurea (HU) treatment, which causes stalled DNA replication forks due to depletion of dNTP pools **(Figure 4E).** Importantly, we observed no sensitivity of USP11 knockdown cells compared with the negative control to treatment with mitomycin c (MMC), which generates interstrand DNA crosslinks **(Figure 4F)**. Similarly, USP11 knockdown in A549 cells sensitized cells to CPT **(Supplemental Figure 2B)**, VP16 **(Supplemental Figure 2C)**, and formaldehyde (**Supplemental Figure 2D**) treatment. USP11 knockdown A549 cells displayed mild sensitivity to HU (**Supplemental Figure 2E**) and no sensitivity to MMC (**Supplemental Figure 2F**) treatment. These observations suggest that USP11 deubiquitinase is required for cell survival upon treatment with DNA-protein, but not DNA-DNA crosslinking agents.

**Figure 4.**
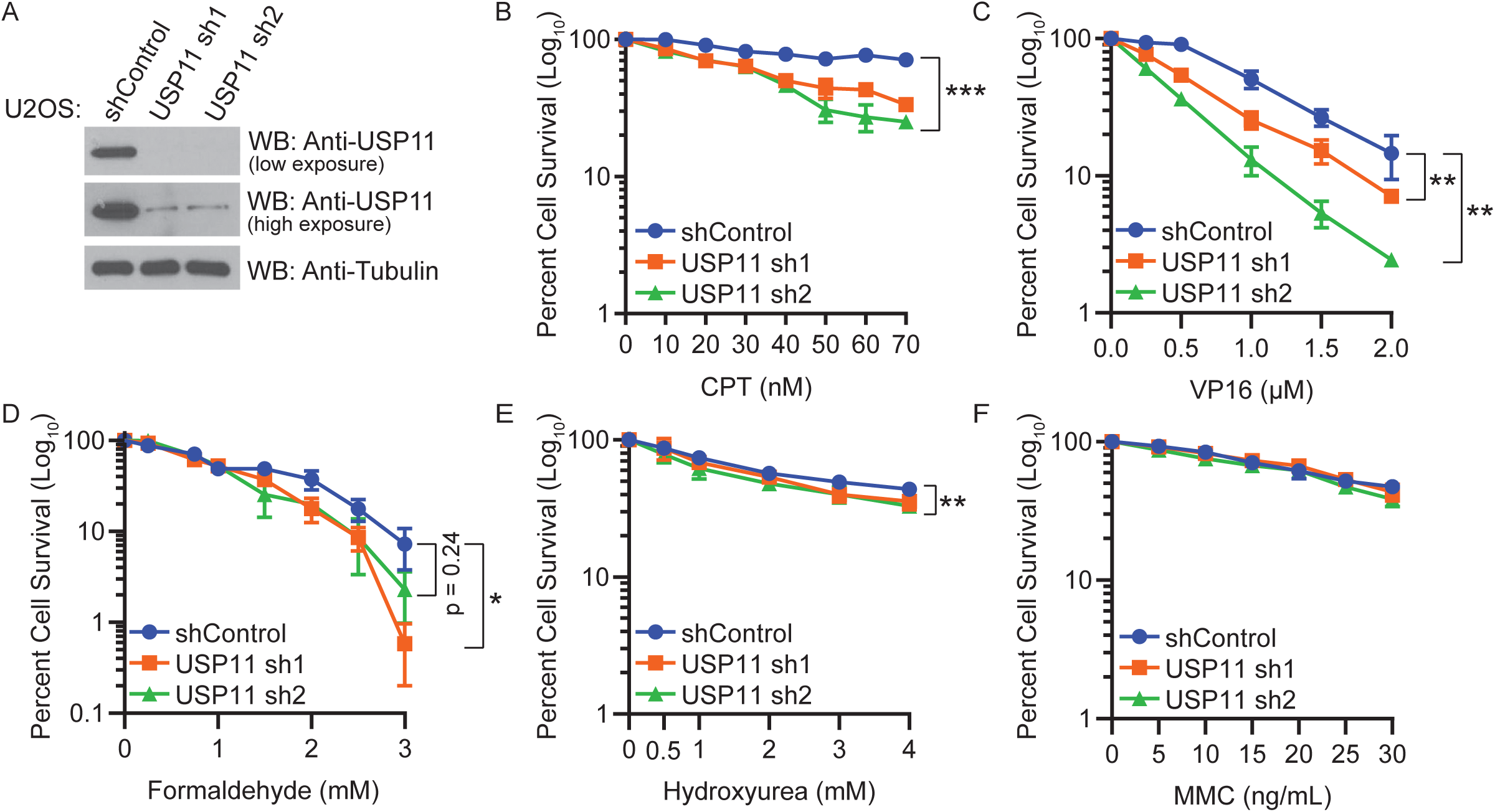
USP11 is required for cell survival upon DNA damage by DPC-inducing agents. (A) WB showing knockdown efficiency of USP11 in U2OS cells stably expressing non-silencing shRNA control or two different shRNA sequences (sh1 and sh2) targeted to USP11. (B-F) Clonogenic cell survival curves for shRNA control and USP11 knockdown U2OS treated with indicated concentrations of (B) CPT for 24 hr, (C) VP16 for 4 hr, (D) formaldehyde for 20 min, (E) HU for 24 hr, or (F) MMC for 24 hr. Percent cell survival was calculated. Data are presented as mean ± SD (n = 3). Statistical analysis: Two-tailed paired t-test was performed using confidence interval = 90% and α = 0.1; *p ≤ 0.1, **p ≤ 0.05, ***p ≤ 0.01. WB, Western Blot. See also Supplemental Figure 2.

### USP11 deubiquitinates SPRTN upon DPC induction

Our data shows that SPRTN is a substrate of USP11 deubiquitinase (**Figure 3**). To examine the effect of USP11 on SPRTN protein stability, we expressed SFB-SPRTN E112A in combination with increasing concentrations of either Myc-USP11 or Myc-USP11 C318S mutant. SPRTN E112A mutant was used to prevent changes in SPRTN protein levels mediated by auto-proteolysis. Increasing concentrations of USP11, but not USP11 C318S, led to a reduction in monoubiquitinated SPRTN **(Supplemental Figure 3A)**. Importantly, SPRTN levels remained unchanged with increasing concentration of USP11 C318S deubiquitinase inactive mutant, suggesting that USP11 does not regulate the stability of SPRTN **(Supplemental Figure 3A)**. Further, we observed a decrease in endogenous unmodified SPRTN levels with a subsequent increase in Ub-SPRTN in USP11 knockdown cells compared with negative control (**Supplemental Figure 3B**), indicating that USP11 deubiquitinates monoubiquitinated SPRTN and does not regulate SPRTN protein stability.

A previous study showed that upon DNA damage by DPC-inducing agents, monoubiquitinated SPRTN is deubiquitinated and localized to chromatin (Stingele et al., 2016). We examined deubiquitination of SFB-SPRTN and SFB-SPRTN E112A catalytic mutant in HEK 293T cells upon treatment with formaldehyde. We observed a stepwise reduction in monoubiquitinated SPRTN FL **(Supplemental Figure 4A)** and SPRTN E112A **(Supplemental Figure 4B)** levels with increasing concentrations of formaldehyde. We next asked if USP11 is required for SPRTN deubiquitination in response to DPCs. To address this question, we introduced SFB-SPRTN into negative control and USP11 knockdown HEK 293T **(Figure 5A)** and HCT116 cell lines **(Figure 5B)** and observed monoubiquitinated SPRTN levels in the absence and presence of DPCs generated by formaldehyde treatment. Monoubiquitinated SPRTN levels were increased in the absence of USP11 compared with the negative control in both HEK 293T **(Figure 5A, lanes 1, 3 and 5)** and HCT116 **(Figure 5B, lanes 1 and 3)** untreated cells. Importantly, we observed a reduction in SFB-SPRTN deubiquitination in USP11 knockdown HEK 293T (**Figure 5A, lanes 4 and 6**) compared with the negative control (**Figure 5A, lane 2**) upon formaldehyde treatment. Similarly, USP11 depletion in HCT116 inhibited SFB-SPRTN deubiquitination (**Figure 5B, lane 4**) compared with negative control **(Figure 5B, lane 2)** upon formaldehyde treatment, suggesting that USP11 depletion inhibits SPRTN deubiquitination upon DPC induction. We next examined deubiquitination of endogenous SPRTN in the absence and presence of DPCs in USP11 knockdown HCT116 and A549 cells. USP11 depletion led to an increase in monoubiquitinated SPRTN levels compared with the negative control in the absence of damage in HCT116 **(Figure 5C, lanes 1 and 3)** and A549 (**Figure 5D, lanes 1 and 3**) cells. Notably, SPRTN deubiquitination was abrogated in USP11 knockdown HCT116 **(Figure 5C, lane 4)** and A549 **(Figure 5D, lane 4)** cells upon treatment with formaldehyde. Collectively, these findings demonstrate that USP11 is required for SPRTN deubiquitination upon DPC induction.

**Figure 5.**
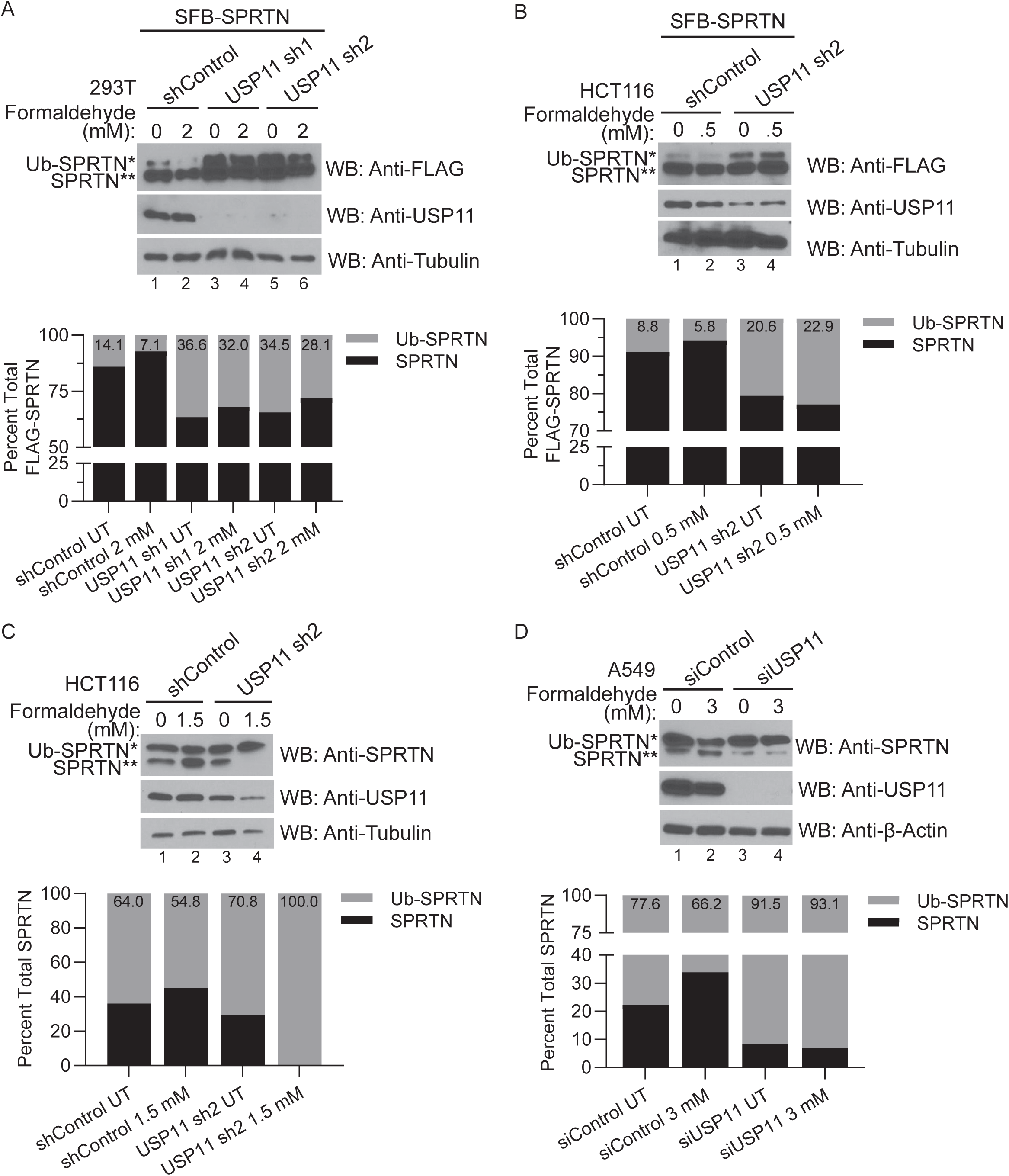
USP11 deubiquitinates SPRTN upon DPC induction. (A-B) HEK 293T (A) or HCT116 (B) cells stably expressing shRNA targeted to USP11 were transfected with SFB-SPRTN. 24 hr later, cells were treated with the indicated concentrations of formaldehyde for 2 hr. (C) HCT116 cells stably expressing shRNA targeted to USP11 were treated with the indicated concentrations of formaldehyde for 1 hr. The nuclear fraction was isolated and used for WB. (D) A549 cells were transfected with siRNA targeted to USP11. 72 hr later, cells were treated with the indicated concentrations of formaldehyde for 2 hr. (A-D) Top panel, cell lysates were immunoblotted with the indicated antibodies. Bottom panel, graph shows percent total FLAG-SPRTN or percent total SPRTN quantified from anti-FLAG or anti-SPRTN blots, respectively. The percent of unmodified and monoubiquitinated SPRTN was calculated. WB, Western Blot. See also Supplemental Figure 3 and Supplemental Figure 4.

### USP11 participates in DPC repair

Upon DPC induction, SPRTN is deubiquitinated and localized to chromatin for DPC repair (Stingele et al., 2016). Our data shows that USP11 is required for SPRTN deubiquitination (**Figure 5**) and cell survival (**Figure 4**) upon DPC induction, indicating that USP11 is required for DPC repair. To demonstrate that USP11 participates in DPC repair, we examined the accumulation and repair of specific DPCs in USP11 knockdown cells using a Rapid Approach to DNA Adduct Recovery (RADAR) assay. We examined TOP1-cc levels in negative control and USP11 knockdown A549 cells and found that USP11 depletion led to an increase in TOP1-ccs in the absence of DPCs induced by treatment with exogenous DPC-inducing agents (**Figure 6A**). We next treated the negative control and USP11 knockdown A549 cells with CPT and examined TOP1-cc induction. With increasing concentration of CPT treatment, USP11 knockdown cells showed an accumulation of TOP1-ccs compared with the negative control **(Figure 6B)**. We further corroborated this finding in USP11 knockdown U2OS cells (**Figure 6C and Supplemental Figure 5A**). We next examined DPC repair by allowing the cells to recover and repair TOP1-ccs generated from CPT treatment. We found that depletion of USP11 led to delayed TOP1-cc repair, indicated by sustained TOP1-ccs following 30 min and 2 hr recovery time periods, while the majority of TOP1-ccs generated in negative control cells were repaired within 30 min **(Figure 6D)** or 45 min **(Supplemental Figure 5B)** and repair was completed by the 2 hr recovery time point **(Figure 6D and Supplemental Figure 5B)**. These findings demonstrate that USP11 is required for the repair of DPCs.

**Figure 6.**
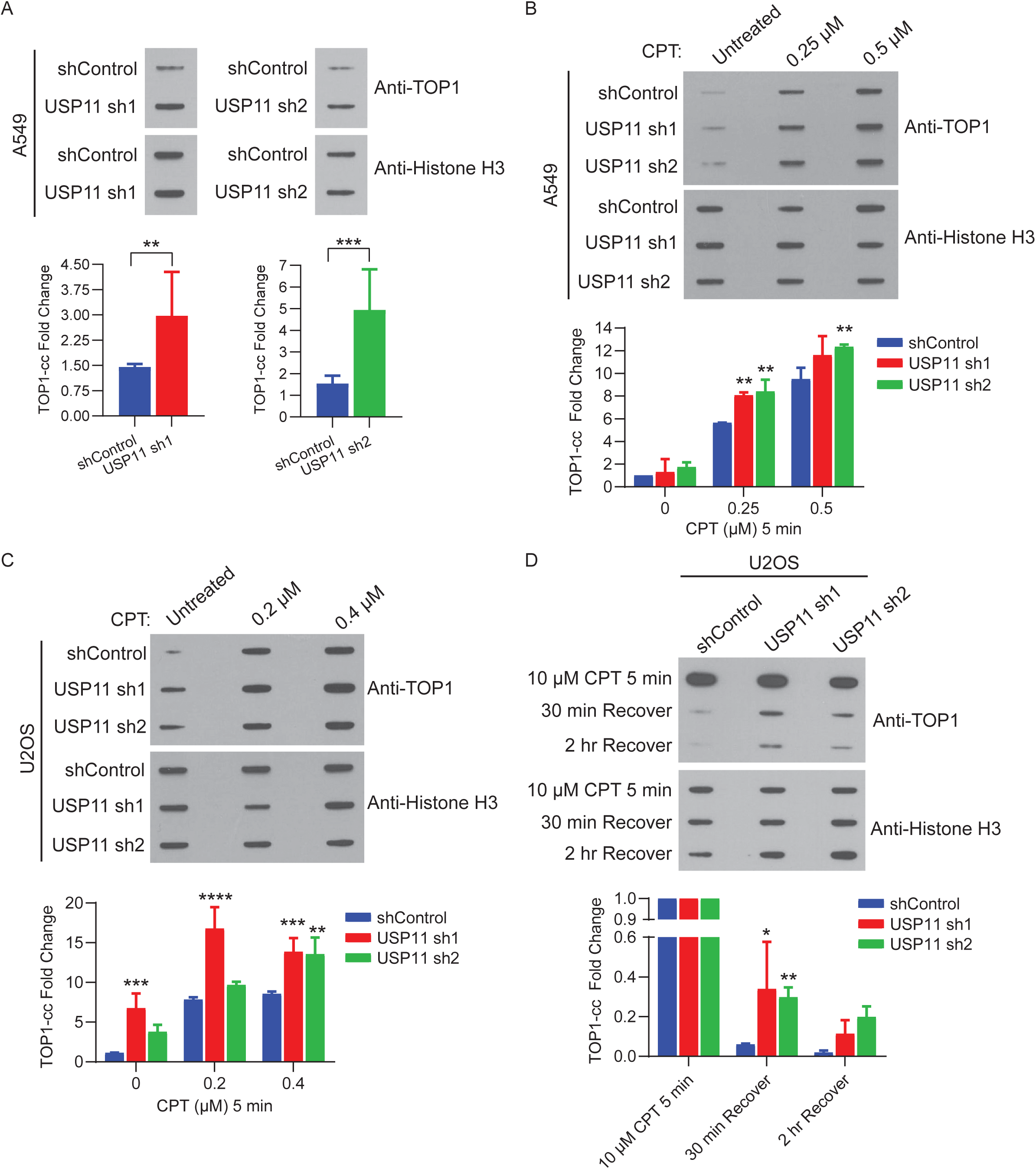
USP11 participates in DPC repair. (A-D) RADAR assays in cells stably expressing non-silencing shRNA control or two different shRNA sequences (sh1 and sh2) targeted to USP11. Sample preparation and slot blotting was completed as described in Materials and methods. Top panel, (A) 500 ng of DNA isolated from untreated A549 cells was immunoblotted with TOP1 antibody to detect endogenous TOP1-ccs and histone H3 was immunoblotted as a loading control. (B) A549 cells were treated with CPT at the indicated concentrations for 5 min. 1 μg of DNA was immunoblotted with TOP1 antibody and 1 μg of DNA was immunoblotted with histone H3. (C) U2OS cells were treated with CPT at the indicated concentrations for 5 min. 1 μg of DNA was immunoblotted with TOP1 antibody and 500 ng of sample was immunoblotted with histone H3. (D) U2OS cells were treated with 10 μM CPT for 30 min. Cells used for recovery time points were washed with PBS and left to recover in drug-free media for the indicated time. 600 ng of DNA was immunoblotted with TOP1 antibody and with histone H3. (A-D) Bottom panel, TOP1-cc fold change from respective slot blots shown above was quantified from relative abundance of TOP1 normalized to histone H3 (A, n = 6; C, n = 2) or normalized to amount of DNA loaded (B, n =2; D, n = 2). Data are presented as mean ± SD. Statistical analysis: Two-way ANOVA and Dunnett’s multiple comparison test was performed using confidence interval = 90% and α = 0.1; *p ≤ 0.1, **p ≤ 0.05, ***p ≤ 0.01, ****p ≤ 0.001. Supplemental Figure 5.

## DISCUSSION

In this study, we showed that USP11 interacts with SPRTN protease and cleaves monoubiquitinated SPRTN to repair DPCs. Based on our findings and previous observations made by others, we propose a model for the regulation of monoubiquitinated SPRTN and recruitment of SPRTN to chromatin during DPC repair **(Figure 7)**. SPRTN protein levels peak during S-phase (Mosbech et al., 2012) and an unknown E3 ligase (X) monoubiquitinates SPRTN, which excludes SPRTN from chromatin (Stingele et al., 2016). Upon DPC induction, such as TOP1-ccs generated by CPT treatment, the DNA replication machinery stalls. USP11 interacts with the SprT domain of SPRTN **(Figure 2A and 2D)** and cleaves the monoubiquitin on SPRTN **(Figure 3 and 5)**. Deubiquitinated SPRTN is recruited to the DPC lesions on chromatin (Stingele et al., 2016). Binding to ssDNA at the DPC site activates SPRTN protease activity, and SPRTN proteolyses the DPC into a peptide-DNA adduct (Stingele et al., 2016, Vaz et al., 2016, Larsen et al., 2019, Li et al., 2019). Following DPC proteolysis, the resulting peptide-DNA substrate is either bypassed by TLS or processed by TDP1/TDP2 to remove the peptide crosslinked to DNA, generating DNA breaks that are repaired by SSB or DSB repair pathways (Stingele et al., 2017).

**Figure 7.**
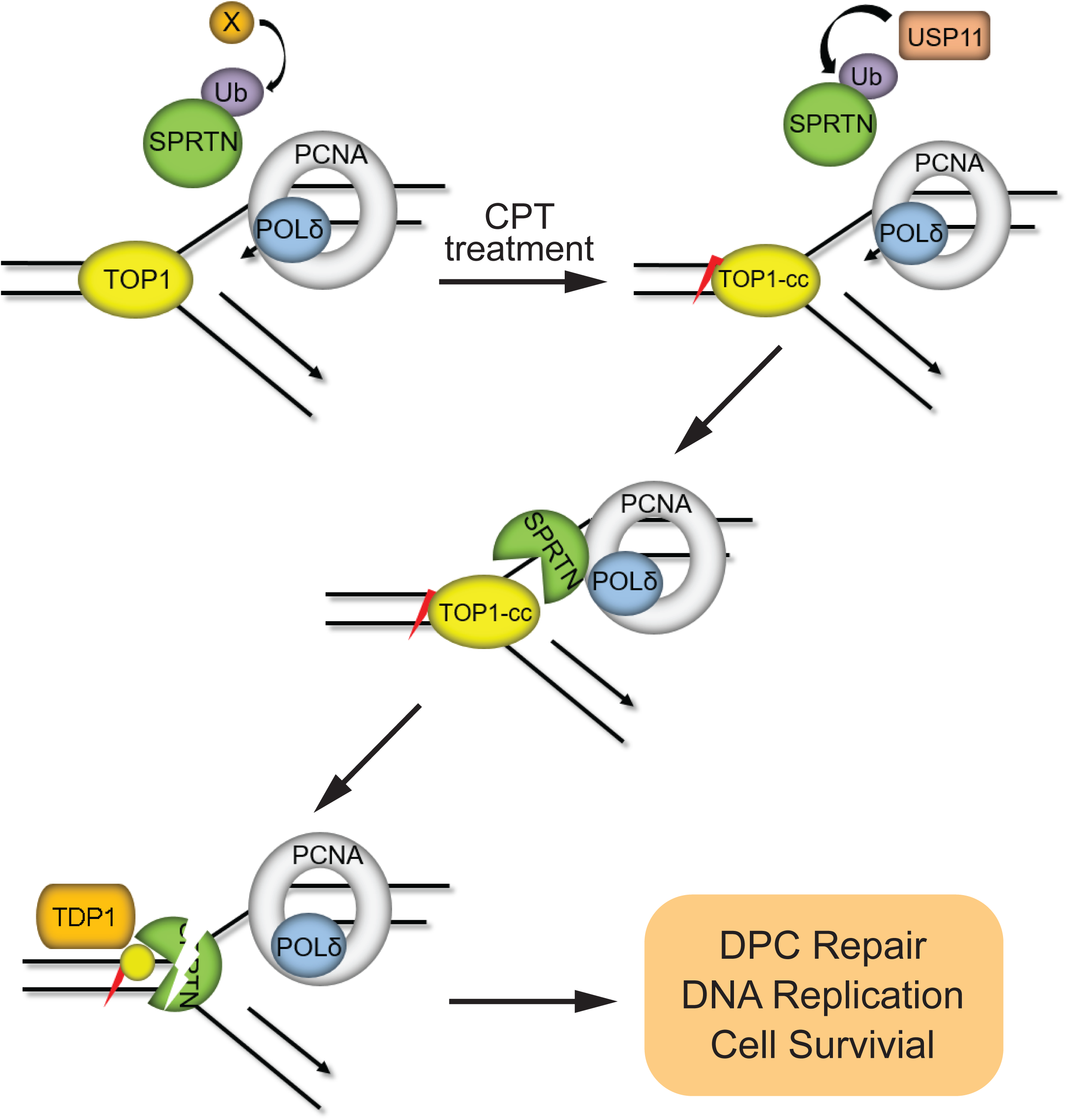
Model for regulation of SPRTN by USP11 in DPC repair.

SPRTN-mediated DPC repair and regulation of SPRTN function is only beginning to be understood. A previous study showed that monoubiquitination of SPRTN is essential for regulation of SPRTN localization and function in response to DPC. However, mutational analysis indicated that the actual site of ubiquitin modification in the C-terminus of SPRTN is not critical, as the ubiquitin can be transferred to an alternative lysine residue with no impact on SPRTN function (Stingele et al., 2016). The E3 ligase that monoubiquitinates SPRTN is not known. We have identified USP11 as a SPRTN ubiquitin protease that deubiquitinates monoubiquitinated SPRTN upon treatment with DPC-inducing agents **(Figure 5)**. However, we observed that depletion of USP11 did not abolish SPRTN deubiquitination **(Figure 5A-D)**. This could be due to incomplete knockdown of USP11 or that one or more DUBs may regulate SPRTN deubiquitination in response to DPCs. Given that SPRTN is a sequence non-specific metalloprotease, promotes the repair of enzymatic and non-enzymatic DPCs, and participates in proteasome-dependent and -independent DPC repair processes, we predict that the tight regulation of SPRTN recruitment to different types of DPC lesions may be regulated by multiple DUBs that are activated by specific DPC-inducing agents to prevent aberrant SPRTN protease activity on chromatin-bound proteins.

Although we observed a slight increase in USP11-SPRTN interaction upon DNA damage with a DPC-inducing agent (**Figure 1D**), it is not clear what prevents SPRTN deubiquitination by USP11 in the absence of damage. A possible factor could be regulated recruitment of USP11 to SPRTN governed by DPC sensors and DNA repair effector proteins. Alternatively, the interaction of USP11 with SPRTN and activation of USP11 may be regulated by the post-translational modifications on USP11 and SPRTN. A recent study showed that SPRTN is phosphorylated at the C-terminus by CHK1 kinase and this phosphorylation enhanced SPRTN recruitment to chromatin (Halder et al., 2019). The cross-talk between ubiquitination-deubiquitination and phosphorylation on SPRTN protease activity and recruitment to DPC lesions remains to be examined. Future studies will involve the identification of DPC sensors/ effectors, deubiquitinases activated by specific types of DPC-lesion and an in-depth analysis on the effect of post-translational modifications of SPRTN and USP11 on USP11 recruitment and function in SPRTN-mediated DPC repair. Further, the identification of the E3 ligase for SPRTN will be critical in deciphering the dynamic regulation of SPRTN ubiquitination and function in DPC repair.

## ACKNOWLEDGEMENTS

This work was supported by NIGMS COBRE (P20 GM121316; G.G. as project leader) and NCI K22 Career Development Award (K22CA188181) to G.G. We thank all members of the Ghosal lab for advice and technical assistance. We thank Dr. Junjie Chen for Gateway cloning vectors. We thank Ross Tomaino at the Taplin Biological Mass Spectrometry Facility at Harvard University for MS data analysis. We thank Dr. Polina Shcherbakova and Dr. Robert Lewis for helpful discussions on the project and critical reading of the manuscript.

## AUTHOR CONTRIBUTIONS

Conceptualization, G.G. and M.P.; Methodology, G.G. and M.P.; Investigation, M.P., S.S.K., M.B., G.S., and M.K.; Resources, G.G., M.P., S.S.K., H.M., N.K., A.H., and M.B.; Writing - Original Draft, G.G. and M.P.; Supervision, G.G.; Funding Acquisition, G.G.

## DECLARATION OF COMPETING INTERESTS

The authors declare no competing interests.

## SUPPLEMENT FIGURE LEGENDS

**Supplemental Figure 1.**
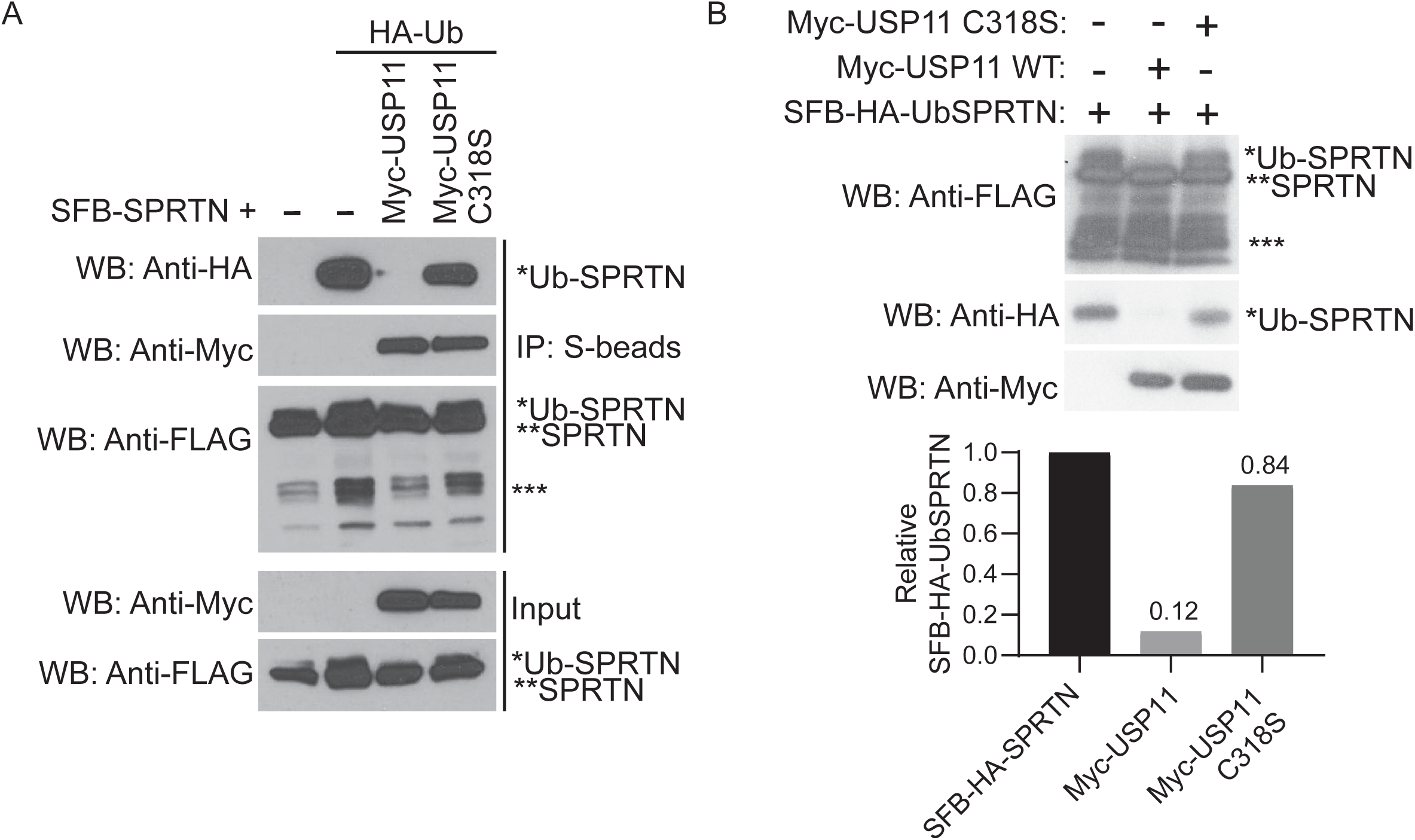
USP11 deubiquitinates SFB-HA-UbSPRTN. (A) HEK 293T cells were transfected with SFB-SPRTN, HA-Ubiquitin and Myc-USP11 FL or C318S. Cell lysates were immunoprecipitated with S-protein agarose beads and immunoblotted with indicated antibodies. (B) Top panel, SFB-HA-UbSPRTN, Myc-USP11, and Myc-USP11 C318S were purified as described in methods. SFB-HA-UbSPRTN was incubated with Myc-USP11 or Myc-USP11 C318S mutant proteins overnight at 30 °C. The reaction mixture was immunoblotted with indicated antibodies. Bottom panel, quantification of anti-HA WB, values relative to SFB-SPRTN alone were calculated and plotted as Relative SFB-HA-UbSPRTN. WB, Western Blot. ***indicates SPRTN auto-cleavage products.

**Supplemental Figure 2.**
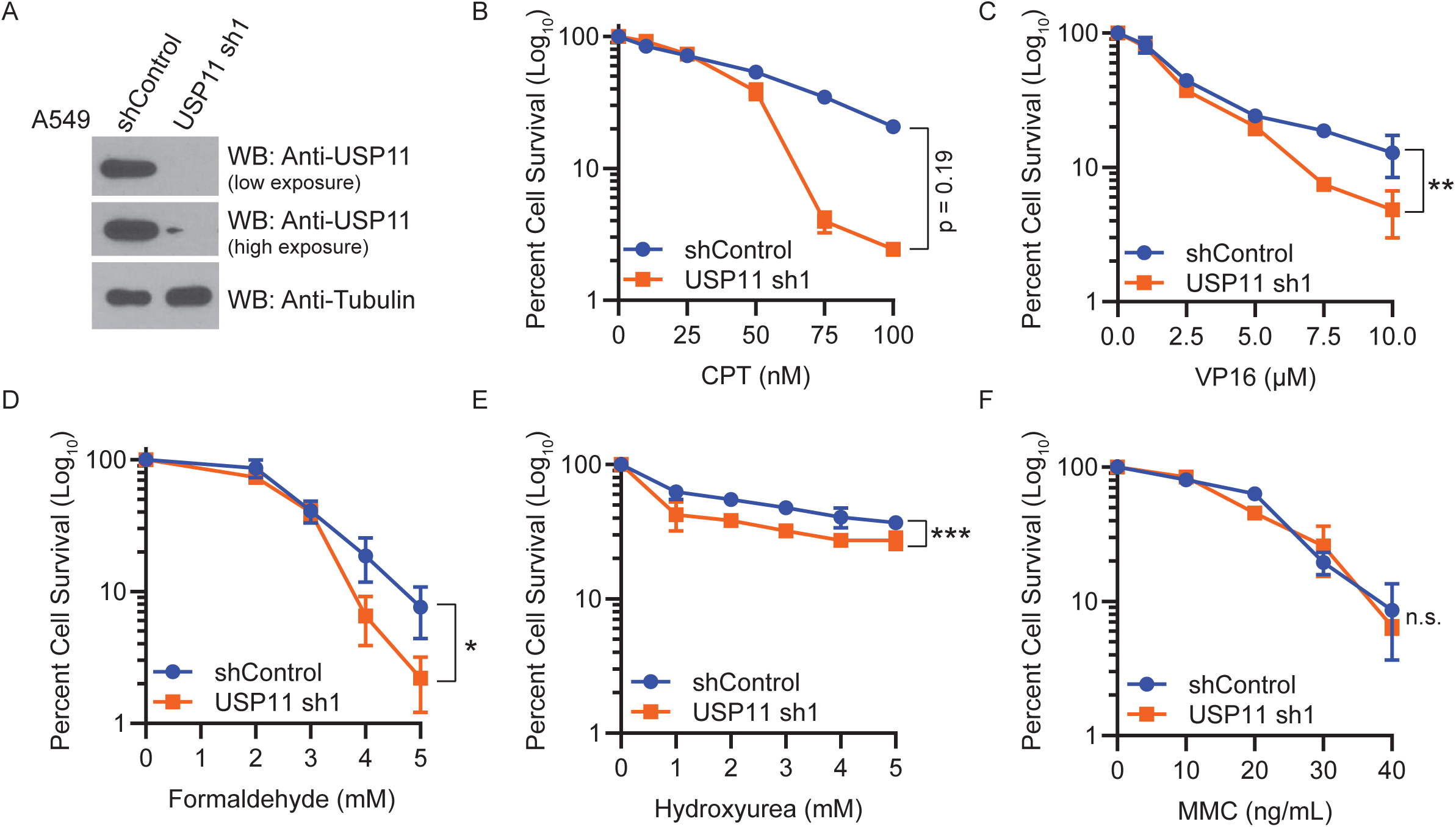
Depletion of USP11 sensitizes cells to treatment with DPC-inducing agents. (A) WB showing knockdown efficiency of USP11 in A549 cells stably expressing non-silencing shRNA control or shRNA targeted to USP11. (B-F) Clonogenic cell survival curves for shRNA control and USP11 knockdown cells treated with indicated concentrations of (B) CPT for 24 hr, (C) VP16 for 4 hr, (D) formaldehyde for 20 min, (E) HU for 24 hr, or (F) MMC for 24 hr. Percent cell survival was calculated and data are presented as mean ± SD (n = 3). Statistical analysis: Two-tailed paired t-test was performed using confidence interval = 90% and α=0.1; *p ≤ 0.1, **p ≤ 0.05, ***p ≤ 0.01; n.s., not statistically significant. WB, Western Blot.

**Supplemental Figure 3.**
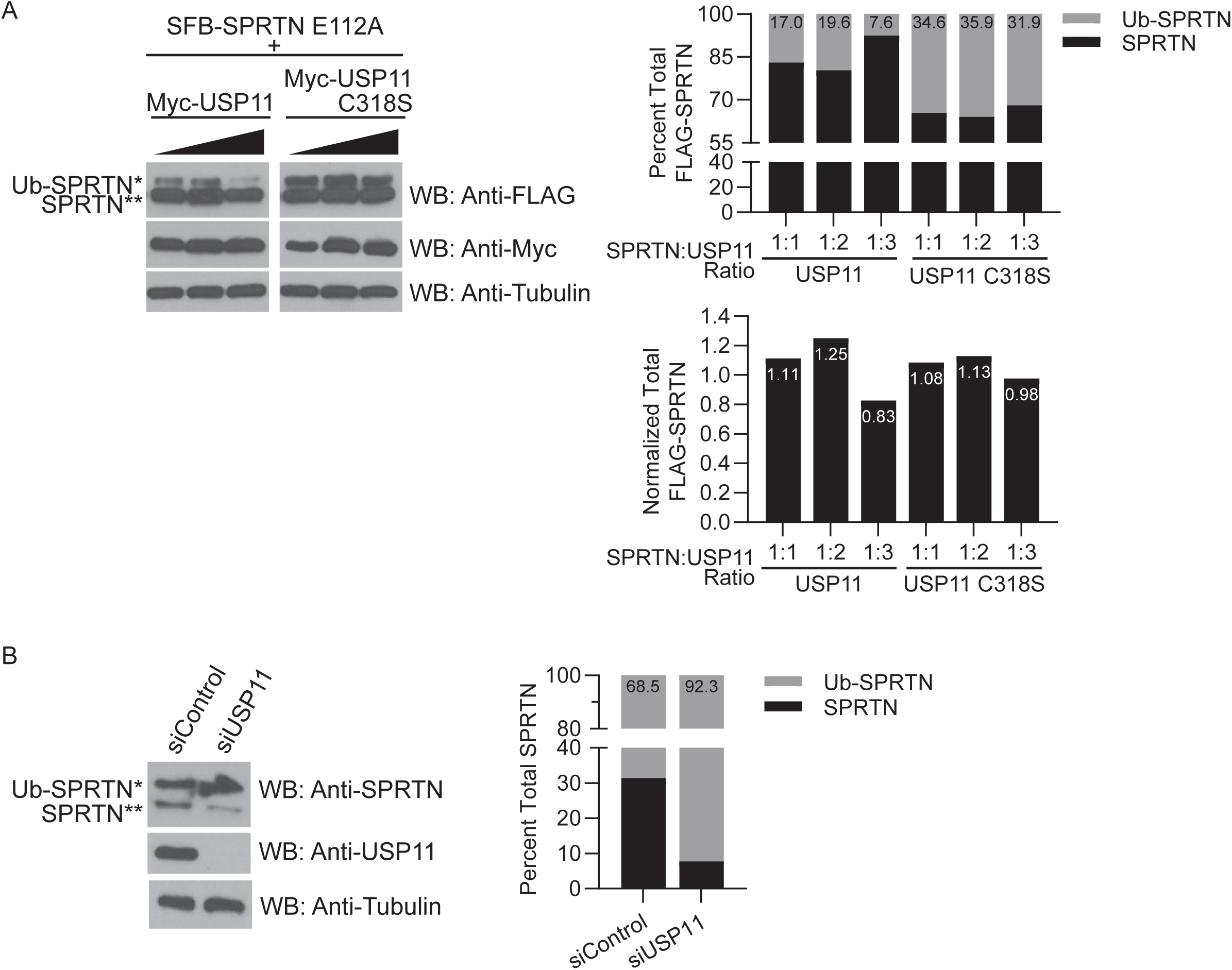
USP11 does not regulate SPRTN protein stability. (A) Left panel, HEK 293T cells were transfected with SFB-SPRTN E112A (2.5 μg) and increasing concentrations (2.5, 5.0, or 7.5 μg) of Myc-USP11 FL or C318S mutant. Cell lysates were immunoblotted with indicated antibodies. Right top panel, quantification of anti-FLAG blots. The percent of unmodified and monoubiquitinated SPRTN was calculated. Graph shows percent total FLAG-SPRTN. Right bottom panel, quantification of anti-FLAG blots. Values of total SFB-SPRTN relative to 1:1 SPRTN:USP11 FL or C318S lanes were calculated and normalized with anti-Tubulin blots and plotted as Normalized total FLAG-SPRTN. (B) Left panel, A549 cells were transfected with siRNA targeted to USP11. 72 hr later, cell lysates were immunoblotted with the indicated antibodies. Right panel, quantification of anti-SPRTN blot. The percent of unmodified and monoubiquitinated SPRTN was calculated and plotted as percent total SPRTN.

**Supplemental Figure 4.**
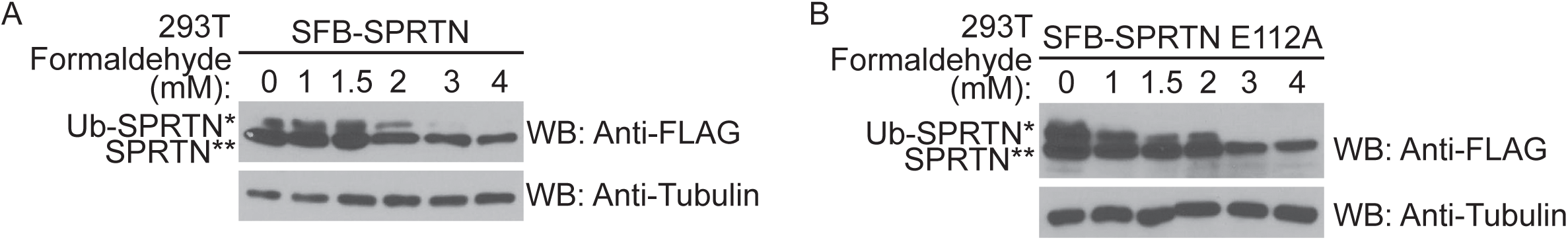
SPRTN is deubiquitinated upon treatment with formaldehyde. HEK 293T cells (A) transfected with SFB-SPRTN or (B) stably expressing SFB-SPRTN E112A were treated with the indicated concentrations of formaldehyde for 2 hr. WB, Western Blot.

**Supplemental Figure 5.**
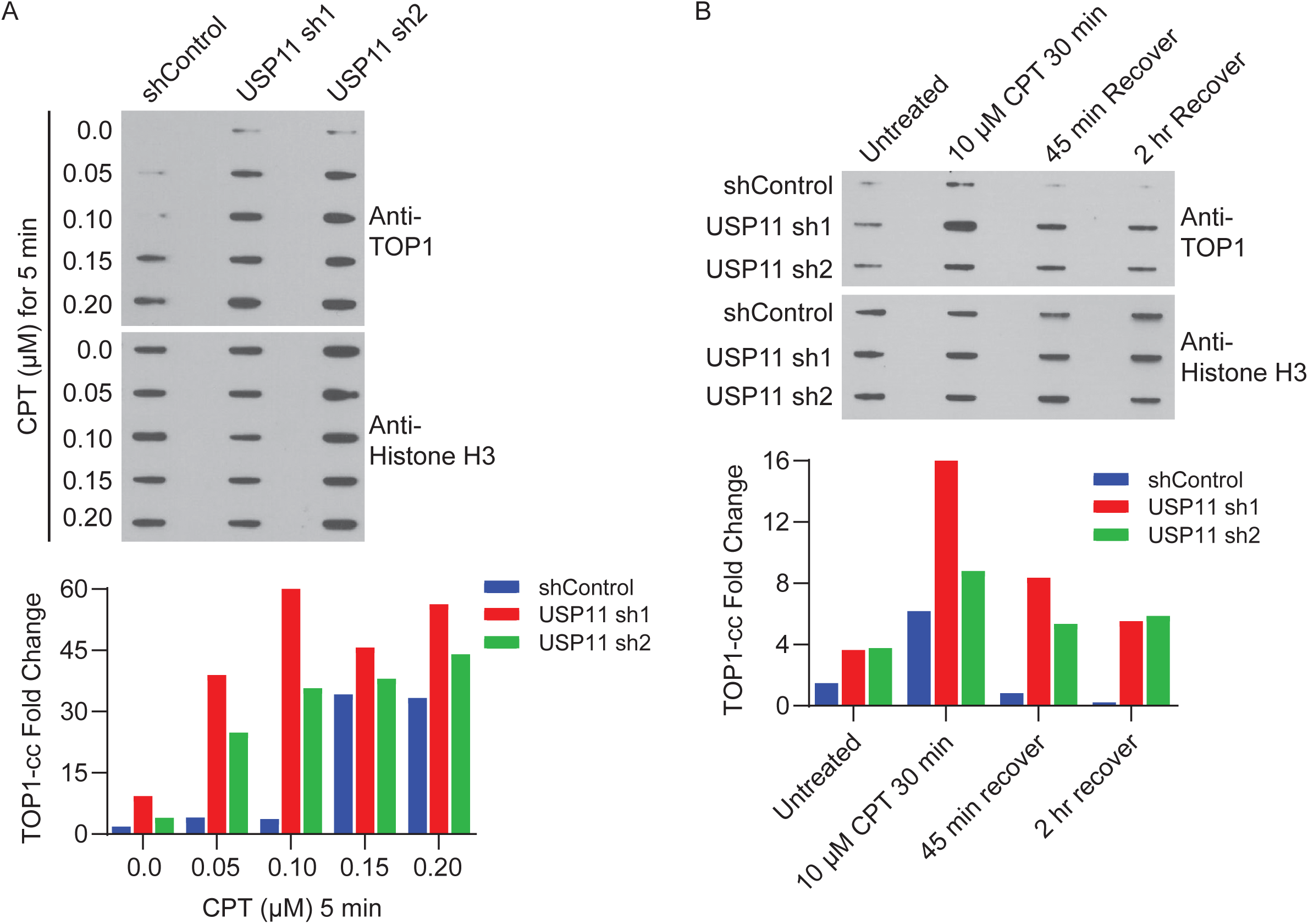
USP11 depleted cells accumulate TOP1-ccs. (A-B) RADAR assays in U2OS cells stably expressing non-silencing shRNA control or two different shRNA sequences (sh1 and sh2) targeted to USP11. Top panel, (A) cells were treated with CPT at the indicated concentrations for 5 min. 300 ng of DNA was immunoblotted with TOP1 antibody and 600 ng of DNA was immunoblotted with histone H3. (B) Cells were treated with 10 μM CPT for 30 min. Cells used for recovery time points were washed with PBS and left to recover in drug-free media for the indicated time. 300 ng of DNA was immunoblotted with TOP1 antibody and with histone H3. Bottom panel, (A-B) TOP1-cc fold change from slot blots shown above was quantified from relative abundance of TOP1 normalized to histone H3.

## SUPPLEMENTAL FILES

**Supplemental File 1:** SPRTN MS analysis results

**Supplemental File 2:** USP11 MS analysis results

**Supplemental File 3:** Oligonucleotides and Primers used in this study

## MATERIALS and METHODS

### Cell culture and Transfections

HEK 293T, U2OS, and A549 cell lines (ATCC) were cultured in Dulbecco’s Modified Eagle medium (DMEM, Hyclone) supplemented with 10% fetal bovine serum (FBS, Hyclone) and 1% penicillin/streptomycin (Gibco). HCT116 cell line (ATCC) was cultured in McCoy’s 5A supplemented with 10% FBS and 1% penicillin/streptomycin. Cells were maintained in a 37°C incubator containing 5% CO_2_. Transfection of expression vectors was carried out using Lipofectamine 2000 (Invitrogen) or PEI (Polysciences, Inc.) reagents as per the protocol from the manufacturer. Non-targeting control siRNA and siRNA targeting USP11 were transfected into cells using Lipofectamine RNAi MAX (Invitrogen) following the manufacturer’s protocol.

### Plasmid cloning and Site-directed mutagenesis

All cDNAs were subcloned into pDONR201 vector as entry clones and N-terminal tagged fusion constructs were generated by transferring the gene insert from the entry clones into gateway-compatible destination vectors by gateway cloning strategy using BP clonase and LR clonase enzymes (ThermoFisher Scientific) as per the manufacturer’s instructions. All deletion and point mutants were generated by Quick change site-directed mutagenesis protocol using KOD Hot Start Polymerase (Millipore) and DpnI (New England Biolabs) digestion. All constructs generated in this study were confirmed by DNA sequencing. See Supplemental File 3 for a complete list of primers used for site-directed mutagenesis.

### Stable cell line generation

Stable shRNA knockdown cell lines were generated by lentiviral transduction of shRNA targeted to USP11 or non-silencing control. Virus was produced in HEK 293T cells transfected with the shRNA plasmid, psPAX2 and pMD2.G lentiviral packaging plasmids in a ratio of 4:2:1. 48 hr post-transfection, virus was collected and used to transduce U2OS, A549, HEK 293T or HCT116 cells. GFP positive cells were sorted by flow cytometry into 96 well plates, and stable clones were selected with puromycin-containing media. See Supplemental File 3 for shRNA sequences. Stable overexpression cell lines were generated by transfecting HEK 293T with SFB-tagged constructs and seeding cells at low density in puromycin-containing media 24 hr post-transfection. Individual clones were isolated and screened for expression. Puromycin (Sigma-Aldrich) concentrations used for stable cell lines are as follows: U2OS, 1.2 μg/mL; A549, 3.0 μg/mL; HEK 293T, 1.0 μg/mL; HCT116 0.25 μg/mL.

### Tandem affinity purification-mass spectrometry

TAP was performed as described previously (Ghosal et al., 2012). HEK 293T cells were transfected with plasmids encoding SFB (S-protein, FLAG, and streptavidin-binding peptide)-tagged constructs. Cell lines stably expressing tagged proteins were selected, and the expression of exogenous proteins was confirmed by immunoblotting and immunostaining. For affinity purification, HEK 293T cells stably expressing SFB-tagged protein from a total of twenty 10-cm dishes were collected and lysed in NETN buffer (250 mM NaCl, 5 mM EDTA pH 8.0, 50 mM Tris-HCl pH 8.0, and 0.5% Nonidet P-40) supplemented with protease inhibitor cocktail (Roche). Crude lysates were cleared by centrifugation, and the supernatants were incubated with 200 μl of streptavidin-Sepharose beads (Sigma-Aldrich) overnight at 4 °C. The beads were washed three times with NETN buffer and then eluted with 2 mg/ml biotin (Sigma-Aldrich) for 2 hr at 4 °C. The eluates were incubated with 40 μl of S-protein-agarose beads (Millipore) for 2 hr at 4 °C and then washed three times with NETN buffer. The proteins bound to beads were eluted by boiling with 4X Laemmli buffer (SDS sample buffer), resolved by SDS-PAGE, visualized by Coomassie Blue staining, and analyzed by mass spectrometry for protein identification performed by the Taplin Biological Mass Spectrometry Facility at Harvard University.

### Immunoblotting

Cells were lysed with NETN buffer on an end-to-end rocker at 4 °C for 30 min. Cleared cell lysates were then collected by centrifugation, boiled in SDS sample buffer, and separated by SDS-PAGE. Proteins were transferred to PVDF membrane (Millipore) via semi-dry transfer. Membranes were blocked in 5% milk in 1X TBS/Tween buffer (TBST) and probed with antibodies as indicated in the figure legends.

### Co-Immunoprecipitations

HEK 293T cells were transfected with constructs encoding SFB and Myc-tagged proteins and incubated for 24 hr. Cells were lysed with NETN buffer. The lysates were cleared by centrifugation and then incubated with 20 μl of S-protein agarose beads overnight on an end-to-end rocker at 4 °C. After three washes with NETN buffer, the proteins bound to beads were eluted by boiling with SDS sample buffer, resolved by SDS-PAGE, and analyzed by Western blotting.

### Clonogenic cell survival assays

800 cells were seeded in 60 mm dishes in triplicate. 24 hr after seeding, cells were treated with CPT (Sigma-Aldrich), VP16 (Sigma-Aldrich), formaldehyde (Sigma-Aldrich), HU (Sigma-Aldrich) or MMC (Sigma-Aldrich) at the indicated concentrations and for the indicated times. Cells were washed with PBS, supplemented with fresh media, and incubated for 10-14 days. Formed colonies were fixed and stained with Coomassie Blue. The numbers of colonies were counted, and the percentage cell survival was calculated.

### *In vitro* deubiquitination assay

The *in vitro* deubiquitination was performed as described (Dupont et al., 2009). SFB-SPRTN alone or HA-Ubiquitin and SFB-SPRTN together were expressed in HEK 293T cells. At 24 hr post-transfection, cells were lysed in NETN buffer. SFB-tagged ubiquitinated SPRTN was purified by immunoprecipitation with streptavidin-sepharose beads followed by elution with 2 mM biotin on an end-to-end rocker at 4 °C for 1hr. The eluate for SFB-HA-UbSPRTN was then immunoprecipitated with anti-HA-beads (ThermoFisher Scientific) and eluted with 2 mg/ml HA peptide (Sigma-Aldrich) at 37 °C for 10 min. In a parallel experiment, Myc-USP11 and Myc-USP11 C318S was expressed in HEK 293T cells for 24 hr. Myc-USP11 or Myc USP11 C318S was purified by immunoprecipitation with anti-Myc-agarose beads (ThermoFisher Scientific) followed by elution with 100 μg/ml c-Myc peptide (Sigma-Aldrich) on an end-to-end rocker at room temperature for 1 hr. For *in vitro* deubiquitination assay, purified SFB-SPRTN or SFB-HA-UbSPRTN was incubated with purified Myc-USP11 or Myc-USP11 C318S in a deubiquitination reaction buffer (50 mM HEPES, pH 7.5, 100 mM NaCl, 5% glycerol, 5 mM MgCl_2_, 1 mM ATP and 1 mM DTT) at 30 °C overnight. The reaction mixture was terminated using SDS-PAGE buffer and analyzed by Western blotting.

### Deubiquitination assay in cells

HEK 293T cells were transfected with constructs encoding SFB-SPRTN, Myc-tagged USP11 FL or C318S and HA-Ubiquitin, where indicated. Cells were lysed 24 hr later with NETN buffer. The lysates were cleared by centrifugation and then incubated with 20 μl of S-protein agarose beads on an end-to-end rocker overnight at 4 °C. After three washes with NETN buffer, the proteins bound to beads were eluted by boiling with SDS sample buffer, resolved by SDS-PAGE, and analyzed by Western blotting.

### Nuclear fraction isolation

Preparation of nuclear fractions was performed as previously described with modifications (Huang et al., 2009). Cell pellets were resuspended in 3 volumes of fractionation lysis buffer (10 mM HEPES pH 7.4, 10 mM KCl, 0.05% NP-40 and protease inhibitors) and incubated at 4 °C for twenty minutes. Nuclei were recovered by centrifugation and resuspended in 2 volumes of low salt fractionation buffer (10 mM Tris-HCl pH 7.4, 0.2 mM MgCl_2_, 1% Triton-X 100 and protease inhibitors). The lysate was incubated at 4 °C for 15 min and the chromatin fraction was pelleted by centrifugation. The supernatant was collected as the nuclear fraction.

### RADAR assay and Slot blotting

Rapid Approach to DNA Adduct Recovery (RADAR) protocol was performed as described (Kiianitsa and Maizels, 2013). 1.5 × 10^5^ cells were seeded in 6-well plates, grown to 80% confluence, and then treated with CPT at the indicated concentrations and time. The cells were directly lysed in 0.5 mL of DNAzol® (Invitrogen). Genomic DNA and DNA-protein covalent complexes were precipitated from the lysate by addition of 0.5 volume of 100% ethanol followed by a 10 min incubation at −20 °C. The precipitate was collected by centrifugation at 7000 rpm for 5 min, washed twice in 70% ethanol, and immediately resuspended in 300 μl of freshly prepared 8 mM NaOH. The DNA content of each sample was measured with a NanoDrop One instrument (ThermoFisher Scientific). Samples were diluted in 25 mM sodium phosphate buffer pH 6.5 to a final volume of 200 μl. Sample was applied to nitrocellulose membrane (Amersham) using a vacuum slot-blot manifold (Hoefer PR648). The membrane was blocked in 5% milk in 1X TBST and incubated with antibodies as described for immunoblotting. TOP1-cc fold change was quantified from relative abundance of TOP1 and normalized to the loading control or to the amount of DNA loaded.

### Quantification and Statistical analysis

All experiments were independently replicated (biological replicates) at least three times. Technical replicates for clonogenic cell survival assays and RADAR assays are indicated in the figure legends. Western blots and slot blots were quantified using ImageJ software and normalized to the loading control or as indicated in the figure legends. Results were graphed and analyzed using GraphPad Prism 8 software and data are presented as mean ± SD. Two-tailed paired t-test was performed for clonogenic cell survival assays and Two-way ANOVA and Dunnett’s multiple comparison test was performed for RADAR assays. All data points were included in determining the p-values with a confidence interval set to 90% and α = 0.1 to account for any variabilities due to data point outliers. Statistical significance was reported as p ≤ 0.1(*), p ≤ 0.05 (**), p ≤ 0.01 (***), p ≤ 0.001 (****) and n.s. as not statistically significant.

### Data and Materials availability

The mass spectrometry proteomics data have been deposited to the ProteomeXchange Consortium via the PRIDE (Perez-Riverol et al., 2019) partner repository with the dataset identifier PXD019923. Further information and requests of materials used in this research should be directed to Gargi Ghosal. Plasmid DNA constructs generated in this study (indicated in the Key Resources Table) will be made available via material transfer agreement (MTA).

| **KEY RESOURCES TABLE** |  |  |
| --- | --- | --- |
| **REAGENT or RESOURCE** | **SOURCE** | **IDENTIFIER** |
| **Antibodies** |  |  |
| Mouse monoclonal anti-FLAG (WB 1:10,000) | Sigma-Aldrich | Cat#F3165<br>RRID:AB_259529 |
| Mouse monoclonal anti-c-Myc (WB 1:1000) | Santa Cruz<br>Biotechnology | Cat#sc-40<br>RRID:AB_627268 |
| Mouse monoclonal anti-β-Actin (WB 1:10,000) | Sigma-Aldrich | Cat#A5441<br>RRID:AB_476744 |
| Mouse monoclonal anti-α-Tubulin (WB 1:5,000) | Sigma-Aldrich | Cat#T6074<br>RRID:AB_477582 |
| Rabbit monoclonal anti-HA-Tag (WB 1:2,000) | Cell Signaling<br>Technology | Cat#3724S<br>RRID:AB_1549585 |
| Mouse monoclonal anti-Ubiquitin (FK2) (WB 1:1,000) | Millipore | Cat#ST1200;<br>RRID: AB_2043482 |
| Rabbit monoclonal anti-K63-linkage Specific Polyubiquitin (WB 1:1,000) | Cell Signaling<br>Technology | Cat#5621S<br>RRID:AB_10827985 |
| Rabbit polyclonal anti-USP11 (WB 1:1000) | Bethyl<br>Laboratories | Cat#A301-613A<br>RRID:AB_1211380 |
| Rabbit polyclonal anti-SPRTN (WB 1:1000) | Invitrogen | Cat#PA5-46262<br>RRID:AB_2605525 |
| Rabbit monoclonal anti-Topoisomerase I (WB 1:2,000) | Abcam | Cat#ab109374<br>RRID:AB_10861978 |
| Rabbit polyclonal anti-Histone H3 (WB 1:2,000) | Cell Signaling<br>Technology | Cat#9715S<br>RRID:AB_331563 |
| Peroxidase-conjugated AffiniPure Polyclonal Goat anti-Rabbit IgG (WB 1:10,000) | Jackson<br>ImmunoResearch<br>Laboratories | Cat#111-035-144;<br>RRID: AB_2307391 |
| Peroxidase-Conjugated AffiniPure Polyclonal Rabbit anti-Mouse IgG + IgM (WB 1:10,000) | Jackson<br>ImmunoResearch<br>Laboratories | Cat#315-035-048;<br>RRID: AB_2340069 |
| **Bacterial Strain** |  |  |
| DH5α Competent Cells | ThermoFisher<br>Scientific | Cat#18265017 |
| **Chemicals, Peptides, Enzymes and Recombinant Proteins** |  |  |
| c-Myc Peptide | Sigma-Aldrich | Cat#M2435 |
| HA Peptide | Sigma-Aldrich | Cat#I2149 |
| DNAzol Reagent® | Invitrogen | Cat#10503027 |
| Etoposide | Sigma-Aldrich | Cat#E1383 |
| (S)-(+)-Camptothecin | Sigma-Aldrich | Cat#C9911 |
| Formaldehyde Solution | Sigma-Aldrich | Cat#F8775 |
| Hydroxyurea | Sigma-Aldrich | Cat#H8627 |
| Mitomycin C | Sigma-Aldrich | Cat#M4287 |
| Pierce Anti-c-Myc Agarose | ThermoFisher<br>Scientific | Cat#20168 |
| Streptavidin Sepharose | Sigma-Aldrich | Cat#GE17-5113-01 |
| S-protein Agarose | Millipore | Cat#69704 |
| Pierce Anti-HA Magnetic Beads | ThermoFisher Scientific | Cat#88836 |
| Biotin | Sigma-Aldrich | Cat#B4501 |
| KOD Hot Start DNA Polymerase | Sigma-Aldrich | Cat#71086 |
| DpnI | New England Biolabs | Cat#R0176L |
| Gateway™ BP Clonase™ Enzyme Mix | ThermoFisher Scientific | Cat#11789021 |
| Gateway™ LR Clonase™ II Enzyme mix | ThermoFisher Scientific | Cat#11791100 |
| Puromycin dihydrochloride from *Streptomyces alboniger* | Sigma-Aldrich | Cat#P7255 |
| **Cell Lines** |  |  |
| Human: HEK 293T/17 | ATCC | Cat#CRL-11268<br>RRID:CVCL_1926 |
| Human: U2OS | ATCC | Cat#HTB-96<br>RRID:CVCL_0042 |
| Human: HCT116 | ATCC | Cat#CCL-247<br>RRID:CVCL_0291 |
| Human: A549 | ATCC | Cat#CCL-185<br>RRID:CVCL_0023 |
| **Recombinant DNA** |  |  |
| Vector: SFB Destination | Dr. Junjie Chen | (Ghosal et al., 2012) |
| Vector: Myc Destination | Dr. Junjie Chen | (Ghosal et al., 2012) |
| Vector: pDNR201 | Dr. Junjie Chen | (Ghosal et al., 2012) |
| Plasmid: HA-Ubiquitin | Addgene | Plasmid#18712<br>RRID:Addgene_18712 |
| Plasmid: SPRTN | Dr. Junjie Chen | (Ghosal et al., 2012) |
| Plasmid: pENTR223-USP11 | Harvard PlasmID | Cat#HsCD00379313 |
| Plasmid: pQFlag-USP7 | Addgene | Plasmid#46751<br>RRID:Addgene_46751 |
| Plasmid: pBluescriptR-SAMHD1 | Harvard Plasmid | Cat#HsCD00333067 |
| Plasmid: psPAX2 lentiviral packaging | Addgene | Plasmid#12260<br>RRID:Addgene_12260 |
| Plasmid: pMD2.G lentiviral packaging | Addgene | Plasmid#12259<br>RRID:Addgene_12259 |
| **Software** |  |  |
| GraphPad Prism 8 | Software | RRID:SCR_002798<br>https://www.graphpad.com/ |
| ImageJ | Software | RRID:SCR_018407<br>https://imagej.nih.gov/ij/ |

## Notes

### Competing Interest Statement

The authors have declared no competing interest.

## REFERENCES

Baker, D. J., Wuenschell, G., Xia, L., Termini, J., Bates, S. E., Riggs, A. D. & O’Connor, T. R. 2007. Nucleotide excision repair eliminates unique DNA-protein cross-links from mammalian cells. J Biol Chem, 282, 22592–604.

Barker, S., Weinfeld, M. & Murray, D. 2005. DNA-protein crosslinks: their induction, repair, and biological consequences. Mutat Res, 589, 111–35.

Centore, R. C., Yazinski, S. A., Tse, A. & Zou, L. 2012. Spartan/C1orf124, a reader of PCNA ubiquitylation and a regulator of UV-induced DNA damage response. Mol Cell, 46, 625–35.

Davis, E. J., Lachaud, C., Appleton, P., Macartney, T. J., Nathke, I. & Rouse, J. 2012. DVC1 (C1orf124) recruits the p97 protein segregase to sites of DNA damage. Nat Struct Mol Biol, 19, 1093–100.

De Graaf, B., Clore, A. & Mccullough, A. K. 2009. Cellular pathways for DNA repair and damage tolerance of formaldehyde-induced DNA-protein crosslinks. DNA Repair (Amst), 8, 1207–14.

Deng, T., Yan, G., Song, X., Xie, L., Zhou, Y., Li, J., Hu, X., Li, Z., Hu, J., Zhang, Y., Zhang, H., Sun, Y., Feng, P., Wei, D., Hu, B., Liu, J., Tan, W. & Ye, M. 2018. Deubiquitylation and stabilization of p21 by USP11 is critical for cell-cycle progression and DNA damage responses. Proc Natl Acad Sci U S A, 115, 4678–4683.

Dupont, S., Mamidi, A., Cordenonsi, M., Montagner, M., Zacchigna, L., Adorno, M., Martello, G., Stinchfield, M. J., Soligo, S., Morsut, L., Inui, M., Moro, S., Modena, N., Argenton, F., Newfeld, S. J. & Piccolo, S. 2009. FAM/USP9x, a deubiquitinating enzyme essential for TGFbeta signaling, controls Smad4 monoubiquitination. Cell, 136, 123–35.

Duxin, J. P., Dewar, J. M., Yardimci, H. & Walter, J. C. 2014. Repair of a DNA-protein crosslink by replication-coupled proteolysis. Cell, 159, 346–57.

Fielden, J., Ruggiano, A., Popovic, M. & Ramadan, K. 2018. DNA protein crosslink proteolysis repair: From yeast to premature ageing and cancer in humans. DNA Repair (Amst), 71, 198–204.

Ghosal, G., Leung, J. W., Nair, B. C., Fong, K. W. & Chen, J. 2012. Proliferating cell nuclear antigen (PCNA)-binding protein C1orf124 is a regulator of translesion synthesis. J Biol Chem, 287, 34225–33.

Halder, S., Torrecilla, I., Burkhalter, M. D., Popovic, M., Fielden, J., Vaz, B., Oehler, J., Pilger, D., Lessel, D., Wiseman, K., Singh, A. N., Vendrell, I., Fischer, R., Philipp, M. & Ramadan, K. 2019. SPRTN protease and checkpoint kinase 1 cross-activation loop safeguards DNA replication. Nat Commun, 10, 3142.

Hu, M., Li, P., Li, M., Li, W., Yao, T., Wu, J. W., Gu, W., Cohen, R. E. & Shi, Y. 2002. Crystal structure of a UBP-family deubiquitinating enzyme in isolation and in complex with ubiquitin aldehyde. Cell, 111, 1041–54.

Huang, J., Huen, M. S., Kim, H., Leung, C. C., Glover, J. N., Yu, X. & Chen, J. 2009. RAD18 transmits DNA damage signalling to elicit homologous recombination repair. Nat Cell Biol, 11, 592–603.

Ide, H., Nakano, T., Salem, A. M. H. & Shoulkamy, M. I. 2018. DNA-protein cross-links: Formidable challenges to maintaining genome integrity. DNA Repair (Amst), 71, 190–197.

Ide, H., Shoulkamy, M. I., Nakano, T., Miyamoto-Matsubara, M. & Salem, A. M. 2011. Repair and biochemical effects of DNA-protein crosslinks. Mutat Res, 711, 113–22.

Jacko, A. M., Nan, L., Li, S., Tan, J., Zhao, J., Kass, D. J. & Zhao, Y. 2016. De-ubiquitinating enzyme, USP11, promotes transforming growth factor beta-1 signaling through stabilization of transforming growth factor beta receptor II. Cell Death Dis, 7, e2474.

Juhasz, S., Balogh, D., Hajdu, I., Burkovics, P., Villamil, M. A., Zhuang, Z. & Haracska, L. 2012. Characterization of human Spartan/C1orf124, an ubiquitin-PCNA interacting regulator of DNA damage tolerance. Nucleic Acids Res, 40, 10795–808.

Kiianitsa, K. & Maizels, N. 2013. A rapid and sensitive assay for DNA-protein covalent complexes in living cells. Nucleic Acids Res, 41, e104.

Kim, M. S., Machida, Y., Vashisht, A. A., Wohlschlegel, J. A., Pang, Y. P. & Machida, Y. J. 2013. Regulation of error-prone translesion synthesis by Spartan/C1orf124. Nucleic Acids Res, 41, 1661–8.

Larsen, N. B., Gao, A. O., Sparks, J. L., Gallina, I., Wu, R. A., Mann, M., Raschle, M., Walter, J. C. & Duxin, J. P. 2019. Replication-Coupled DNA-Protein Crosslink Repair by SPRTN and the Proteasome in Xenopus Egg Extracts. Mol Cell, 73, 574–588 e7.

Lessel, D., Vaz, B., Halder, S., Lockhart, P. J., Marinovic-Terzic, I., Lopez-Mosqueda, J., Philipp, M., Sim, J. C., Smith, K. R., Oehler, J., Cabrera, E., Freire, R., Pope, K., Nahid, A., Norris, F., Leventer, R. J., Delatycki, M. B., Barbi, G., Von Ameln, S., Hogel, J., Degoricija, M., Fertig, R., Burkhalter, M. D., Hofmann, K., Thiele, H., Altmuller, J., Nurnberg, G., Nurnberg, P., Bahlo, M., Martin, G. M., Aalfs, C. M., Oshima, J., Terzic, J., Amor, D. J., Dikic, I., Ramadan, K. & Kubisch, C. 2014. Mutations in SPRTN cause early onset hepatocellular carcinoma, genomic instability and progeroid features. Nat Genet, 46, 1239–44.

Li, F., Raczynska, J. E., Chen, Z. & Yu, H. 2019. Structural Insight into DNA-Dependent Activation of Human Metalloprotease Spartan. Cell Rep, 26, 3336–3346 e4.

Lin, C. H., Chang, H. S. & Yu, W. C. 2008. USP11 stabilizes HPV-16E7 and further modulates the E7 biological activity. J Biol Chem, 283, 15681–8.

Lopez-Mosqueda, J., Maddi, K., Prgomet, S., Kalayil, S., Marinovic-Terzic, I., Terzic, J. & Dikic, I. 2016. SPRTN is a mammalian DNA-binding metalloprotease that resolves DNA-protein crosslinks. Elife, 5.

Luo, Q., Wu, X., Nan, Y., Chang, W., Zhao, P., Zhang, Y., Su, D. & Liu, Z. 2020. TRIM32/USP11 Balances ARID1A Stability and the Oncogenic/Tumor-Suppressive Status of Squamous Cell Carcinoma. Cell Rep, 30, 98–111 e5.

Machida, Y., Kim, M. S. & Machida, Y. J. 2012. Spartan/C1orf124 is important to prevent UV-induced mutagenesis. Cell Cycle, 11, 3395–402.

Maskey, R. S., Flatten, K. S., Sieben, C. J., Peterson, K. L., Baker, D. J., Nam, H. J., Kim, M. S., Smyrk, T. C., Kojima, Y., Machida, Y., Santiago, A., Van Deursen, J. M., Kaufmann, S. H. & Machida, Y. J. 2017. Spartan deficiency causes accumulation of Topoisomerase 1 cleavage complexes and tumorigenesis. Nucleic Acids Res, 45, 4564–4576.

Maskey, R. S., Kim, M. S., Baker, D. J., Childs, B., Malureanu, L. A., Jeganathan, K. B., Machida, Y., Van Deursen, J. M. & Machida, Y. J. 2014. Spartan deficiency causes genomic instability and progeroid phenotypes. Nat Commun, 5, 5744.

Mosbech, A., Gibbs-Seymour, I., Kagias, K., Thorslund, T., Beli, P., Povlsen, L., Nielsen, S. V., Smedegaard, S., Sedgwick, G., Lukas, C., Hartmann-Petersen, R., Lukas, J., Choudhary, C., Pocock, R., Bekker-Jensen, S. & Mailand, N. 2012. DVC1 (C1orf124) is a DNA damage-targeting p97 adaptor that promotes ubiquitin-dependent responses to replication blocks. Nat Struct Mol Biol, 19, 1084–92.

Nakano, T., Katafuchi, A., Matsubara, M., Terato, H., Tsuboi, T., Masuda, T., Tatsumoto, T., Pack, S. P., Makino, K., Croteau, D. L., Van Houten, B., Iijima, K., Tauchi, H. & Ide, H. 2009. Homologous recombination but not nucleotide excision repair plays a pivotal role in tolerance of DNA-protein cross-links in mammalian cells. J Biol Chem, 284, 27065–76.

Nakano, T., Morishita, S., Katafuchi, A., Matsubara, M., Horikawa, Y., Terato, H., Salem, A. M., Izumi, S., Pack, S. P., Makino, K. & Ide, H. 2007. Nucleotide excision repair and homologous recombination systems commit differentially to the repair of DNA-protein crosslinks. Mol Cell, 28, 147–58.

Orthwein, A., Noordermeer, S. M., Wilson, M. D., Landry, S., Enchev, R. I., Sherker, A., Munro, M., Pinder, J., Salsman, J., Dellaire, G., Xia, B., Peter, M. & Durocher, D. 2015. A mechanism for the suppression of homologous recombination in G1 cells. Nature, 528, 422–6.

Perez-Riverol, Y., Csordas, A., Bai, J., Bernal-Llinares, M., Hewapathirana, S., Kundu, D. J., Inuganti, A., Griss, J., Mayer, G., Eisenacher, M., Perez, E., Uszkoreit, J., Pfeuffer, J., Sachsenberg, T., Yilmaz, S., Tiwary, S., Cox, J., Audain, E., Walzer, M., Jarnuczak, A. F., Ternent, T., Brazma, A. & Vizcaino, J. A. 2019. The PRIDE database and related tools and resources in 2019: improving support for quantification data. Nucleic Acids Res, 47, D442–D450.

Pommier, Y., Huang, S. Y., Gao, R., Das, B. B., Murai, J. & Marchand, C. 2014. Tyrosyl-DNA-phosphodiesterases (TDP1 and TDP2). DNA Repair (Amst), 19, 114–29.

Quievryn, G. & Zhitkovich, A. 2000. Loss of DNA-protein crosslinks from formaldehyde-exposed cells occurs through spontaneous hydrolysis and an active repair process linked to proteosome function. Carcinogenesis, 21, 1573–80.

Serbyn, N., Noireterre, A., Bagdiul, I., Plank, M., Michel, A. H., Loewith, R., Kornmann, B. & Stutz, F. 2019. The Aspartic Protease Ddi1 Contributes to DNA-Protein Crosslink Repair in Yeast. Mol Cell.

Stingele, J., Bellelli, R., Alte, F., Hewitt, G., Sarek, G., Maslen, S. L., Tsutakawa, S. E., Borg, A., Kjaer, S., Tainer, J. A., Skehel, J. M., Groll, M. & Boulton, S. J. 2016. Mechanism and Regulation of DNA-Protein Crosslink Repair by the DNA-Dependent Metalloprotease SPRTN. Mol Cell, 64, 688–703.

Stingele, J., Bellelli, R. & Boulton, S. J. 2017. Mechanisms of DNA-protein crosslink repair. Nat Rev Mol Cell Biol, 18, 563–573.

Stingele, J., Schwarz, M. S., Bloemeke, N., Wolf, P. G. & Jentsch, S. 2014. A DNA-dependent protease involved in DNA-protein crosslink repair. Cell, 158, 327–338.

Stockum, A., Snijders, A. P. & Maertens, G. N. 2018. USP11 deubiquitinates RAE1 and plays a key role in bipolar spindle formation. PLoS One, 13, e0190513.

Sun, W., Tan, X., Shi, Y., Xu, G., Mao, R., Gu, X., Fan, Y., Yu, Y., Burlingame, S., Zhang, H., Rednam, S. P., Lu, X., Zhang, T., Fu, S., Cao, G., Qin, J. & Yang, J. 2010. USP11 negatively regulates TNFalpha-induced NF-kappaB activation by targeting on IkappaBalpha. Cell Signal, 22, 386–94.

Ting, X., Xia, L., Yang, J., He, L., Si, W., Shang, Y. & Sun, L. 2019. USP11 acts as a histone deubiquitinase functioning in chromatin reorganization during DNA repair. Nucleic Acids Res, 47, 9721–9740.

Tretyakova, N. Y., Groehler, A. T. & Ji, S. 2015. DNA-Protein Cross-Links: Formation, Structural Identities, and Biological Outcomes. Acc Chem Res, 48, 1631–44.

Tvaz, B., Popovic, M., Newman, J. A., Fielden, J., Aitkenhead, H., Halder, S., Singh, A. N., Vendrell, I., Fischer, R., Torrecilla, I., Drobnitzky, N., Freire, R., Amor, D. J., Lockhart, P. J., Kessler, B. M., Mckenna, G. W., Gileadi, O. & Ramadan, K. 2016. Metalloprotease SPRTN/DVC1 Orchestrates Replication-Coupled DNA-Protein Crosslink Repair. Mol Cell, 64, 704–719.

Wiltshire, T. D., Lovejoy, C. A., Wang, T., Xia, F., O’Connor, M. J. & Cortez, D. 2010. Sensitivity to poly(ADP-ribose) polymerase (PARP) inhibition identifies ubiquitin-specific peptidase 11 (USP11) as a regulator of DNA double-strand break repair. J Biol Chem, 285, 14565–71.

Yu, M., Liu, K., Mao, Z., Luo, J., Gu, W. & Zhao, W. 2016. USP11 Is a Negative Regulator to gammaH2AX Ubiquitylation by RNF8/RNF168. J Biol Chem, 291, 959–67.

