## Supplemental File 3 for "USP11 deubiquitinates monoubiquitinated SPRTN to repair DNA-protein crosslinks"

| **REAGENT or RESOURCE** | SOURCE | IDENTIFIER |
| --- | --- | --- |
| **Oligonucleotides** | | |
| shRNA Targeting Sequence: USP11 #1  CTCATCTTGAAAGAGTGTG | Horizon Discovery | Cat#RHS4531-EG8237  CloneID:V2LHS_41513 |
| shRNA Targeting Sequence: USP11 #2  ACCTCGTCATAGTAACGCT | Horizon Discovery | Cat#RHS4531-EG8237  CloneID:V3LHS_387857 |
| GIPZ Non-silencing Lentiviral shRNA Control | Horizon Discovery | Cat#RHS4346 |
| siRNA Targeting Sequence: USP11  Sense: AAGCGUUACUAUGACGAGGUAUU | Horizon Discovery | (Wiltshire et al., 2010) |
| siRNA Control Targeting Sequence  Sense: AUGAACGUGAAUUGCUCAAUU | Horizon Discovery | Cat#D-001810-01 |
| **Primers** | | |
| SPRTN ΔSprT 5ʹ gtggaccccacaccggacaaggaaccagagaattac 3ʹ | This paper | N/A |
| SPRTN ΔSH 5ʹ gcccagctagtaatccctgaaacaagcaatttacct 3ʹ | This paper | N/A |
| SPRTN ΔPIP 5ʹ gctgttagtaacagtcaccctagagtatcatttgcc 3ʹ | This paper | N/A |
| SPRTN ΔUBZ 5ʹ tcatccagtcagagcaaagaaggtgacagcatcaaa 3ʹ | This paper | N/A |
| SPRTN E112A 5ʹ accctcctgcatgcaatgatacatgcc 3ʹ | This paper | N/A |
| SPRTN Y117C 5ʹ atgatacatgcctgtttatttgtcact 3ʹ | This paper | N/A |
| USP11 C318S 5ʹ ctgggcaacacgtccttcatgaactcg 3ʹ | This paper | N/A |
| USP11 ΔDUSP 5ʹ ccacagcacgaggagctggagctgcccaacatccag 3ʹ | This paper | N/A |
| USP11 ΔUSP  FWD: 5ʹ ggggacaagtttgtacaaaaaagcaggcttcatggcagtagccccgcgactg 3ʹ  REV: 5ʹ ggggaccactttgtacaagaaagctgggtttcagatgcctggctgacc 3ʹ | This paper | N/A |
| USP11 Δ426-452 5ʹ aagaagaaggagtatgtggtgatcgtggacactttc 3ʹ | This paper | N/A |
| USP11 Δ453-479 5ʹ aaacggcggaacgattctcccttctgctacctcagt 3ʹ | This paper | N/A |
| USP11 Δ480-505 5ʹ gtatctgtgaccttcgaccgccgcaagccagagcag 3ʹ | This paper | N/A |
| USP11 Δ651-683 5ʹ aaacccaactcagatgatgctgggcccagctctgga 3ʹ | This paper | N/A |
| USP11 Δ684-716 5ʹ gaccctgagccagagcagttcaccctgcagacggtg 3ʹ | This paper | N/A |
| USP11 Δ717-748 5ʹ cgacgacgcaagcagctgccagagatgaagaagcgt 3ʹ | This paper | N/A |
